# Endosome-escaping engineered LNP-miR146a with in vivo biodistribution to mitigate inflammation and foreign body giant cell formation

**DOI:** 10.64898/2026.02.13.705811

**Authors:** Mohammad I. Khan, Karunakaran R. Sankaran, Shaik O. Rahaman

## Abstract

Nanomaterial-enabled delivery systems have transformed therapeutic strategies for treating inflammatory and degenerative diseases by enabling targeted delivery of small molecules and nucleic acids. However, the clinical translation of microRNA (miR) therapeutics remains limited by instability, enzymatic degradation, and inefficient intracellular delivery in biological environments. Here, we present the design and validation of a next-generation lipid nanoparticle (LNP) platform optimized for the stable and effective delivery of the anti-inflammatory microRNA miR-146a. This LNP system is produced using a scalable lipid injection-based formulation method and yields nanoparticles with uniform size distribution and exceptional physicochemical stability across a wide pH range (2.5-8) and in serum-containing conditions. The four-component lipid architecture enables high miR-146a loading efficiency, efficient endo/lysosomal escape, and robust cellular internalization, resulting in effective tissue uptake and biodistribution both *in vitro* and *in vivo*. Importantly, LNP-mediated delivery of miR-146a exhibits excellent biocompatibility and potent anti-inflammatory activity in primary cells and animal models. Collectively, these results suggest this LNP-miR146a platform as a stable, efficient, and translatable approach for modulating inflammation and addressing biomaterial-associated inflammatory responses.

## Introduction

Nanomaterial-based strategies, including drug-delivery vehicles, wound-dressing systems, and implantable devices, are being actively developed to deliver therapeutic small molecules, proteins, and nucleic acids for the treatment of life-threatening diseases such as cardiovascular disorders, infectious diseases, chronic wounds, inflammatory bowel disease, and autoimmune conditions (1–4). Among these platforms, RNA-loaded LNPs, such as those used in clinically approved vaccines, typically consist of four principal lipid components: ionizable lipids, helper phospholipids, cholesterol, and polyethylene glycol (PEG)-conjugated lipids (5–7). Each component plays a distinct functional role: ionizable lipids acquire a positive charge in acidic environments to facilitate encapsulation of the negatively charged RNA payload; helper phospholipids enhance particle stability and promote membrane fusion; cholesterol modulates membrane integrity and rigidity; and PEGylated lipids confer steric stabilization and prolonged circulation by reducing nonspecific interactions (5–10).

Despite their clinical success, LNP formulations remain vulnerable to chemical degradation (e.g., oxidation and hydrolysis) and enzymatic breakdown in biological fluids, posing significant challenges to long-term stability and shelf life (11,12). To address these limitations and improve miR delivery, we engineered and synthesized a novel LNP formulation composed of four distinct lipid components at a molar ratio of 40:10:48:2. Specifically, the formulation includes the zwitterionic helper phospholipid 1,2-dioleoyl-sn-glycero-3-phosphoethanolamine (DOPE) to promote endosomal escape; the cationic lipid 1,2-dioleoyl-3-trimethylammonium-propane (DOTAP) to enable efficient nucleic acid complexation and delivery; the unsaturated fatty acid linoleic acid to enhance membrane fusion and endosomal escape; and D-α-tocopherol polyethylene glycol 1000 succinate (TPGS) to improve nanoparticle stability and colloidal integrity (8,13).

miRs are small (18-22 nucleotides), single-stranded, non-coding RNAs that bind to the 3’ untranslated region of target mRNAs, leading to post-transcriptional gene silencing through translational repression or mRNA degradation (14–18). Through this mechanism, miRs exert powerful control over gene expression and cellular function. Emerging evidence has established miRs as potent endogenous “master regulators” capable of coordinating complex gene networks that govern numerous functions including cell activation, proliferation, differentiation, inflammation, and fibrosis (14–19). Dysregulated miR expression contributes to the pathogenesis of numerous diseases, including fibrosis, cancer, and the foreign body response (FBR) to implanted biomaterials (20–24). Among anti-inflammatory miRs, miR-146a has emerged as one of the most robust regulators of inflammatory signaling (25–29). miR-146a plays a critical role in immune regulation, cell differentiation, and cell survival by targeting key signaling nodes such as TNF receptor–associated factor 6 (TRAF6), nuclear factor-κB (NF-κB), and Toll-like receptor (TLR) pathways during inflammatory responses (30–38). Our recent studies identify miR-146a not merely as a modulator, but as a central “molecular brake” that limits the severity of the FBR to implanted biomaterials (20). We demonstrated a strict inverse relationship between miR-146a expression and immune activation: elevated miR-146a levels markedly suppress macrophage accumulation, foreign body giant cell (FBGC) formation, and fibrotic encapsulation, whereas genetic deletion of miR-146a in murine models removes this regulatory brake, resulting in exacerbated inflammation and an amplified FBR (20). From a therapeutic perspective, miR-146a delivery to inflamed or injured tissues can be achieved using LNPs, which may protect miRs from nuclease degradation, facilitate endosomal escape, and enhance stability and bioavailability compared to naked miR delivery. miR-loaded LNPs have emerged as versatile and powerful nanocarriers capable of modulating diverse cellular processes, including inflammation, fibrogenesis, cell differentiation, and proliferation (38–42). Thus, LNP-mediated miR-146a delivery holds strong potential as a targeted strategy to modulate host inflammatory responses.

In this study, we report the rational engineering and scalable preparation of a novel miR-146a-encapsulated LNP platform that is uniformly sized and exhibits exceptional stability across a broad pH range (2.5-8) as well as in serum. Using a simple lipid injection and stirring approach, we formulated LNPs composed of four distinct lipid components that enable efficient miR-146a encapsulation, robust endo/lysosomal escape, and effective cellular uptake and biodistribution in vitro and in vivo. Importantly, these LNP-miR146a nanoparticles demonstrate excellent biocompatibility and potent anti-inflammatory activity both in vitro and in vivo, positioning LNP-miR146a delivery as a robust and translatable strategy for controlling host inflammatory responses and mitigating biomaterial-associated inflammation.

## Materials and methods

### Materials

To formulate LNPs, 1,2-dioleoyl-sn-glycero-3-phosphoethanolamine (DOPE 18:1), 1,2-dioleoyl-3-trimethylammonium-propane (chloride salt) (DOTAP 18:1) and 1,2-dioleoyl-sn-glycero-3-phosphoethanolamine-N-(7-nitro-2-1,3-benzoxadiazol-4-yl) (ammonium salt) (NBD PE 18:1) were purchased from Avanti Polar Lipids, USA. Linoleic acid (LA), D-α-Tocopherol polyethylene glycol 1000 succinate (TPGS) and Polyethyleneimine (PEI) were purchased from Millipore-Sigma and all the lipids were used without further purification. MTT (3-[4,5-dimethylthiazol-2-yl]-2,5 diphenyl tetrazolium bromide) assay kit was purchased from Millipore-Sigma. Mouse-recombinant interleukin-4 (IL-4), granulocyte-macrophage colony-stimulating factor (GMCSF), and macrophage colony-stimulating factor (MCSF) were sourced from R&D Systems. We acquired mixed cellulose esters membranes (0.22-μm pore size filters, 12 mm) from Millipore-Sigma. Cell culture essentials like Dulbecco’s modified Eagle’s medium (DMEM), fetal bovine serum (FBS), antibiotic-antimycotic, and related reagents were purchased from Gibco. ProLong Diamond 4′,6-diamidino-2-phenylindole (DAPI) was purchased from Thermo Fisher Scientific.

### LNP formulation

LNPs were formulated using an ethanolic injection method combined with vigorous stirring, as previously described (41). Ethanolic stock solutions of DOPE, DOTAP, TPGS were prepared in 100% ethanol, while LA was used directly in its liquid form. The lipid stocks were further diluted in 100% ethanol to achieve a final molar ratio of 40:10:48:2 for DOPE:DOTAP:LA:TPGS, respectively. The lipids mixture (100 μl) was loaded into a Hamilton syringe and injected into 5 ml of 20 mM HEPES buffer (pH 7.4) under continuous stirring condition and maintained for 60 min. Following stirring at 500 rpm for 60 min, the solution was filtered through a 0.4 μm syringe filter (denoted as 60-F) to remove aggregates and obtain a homogeneous formulation. The final LNP suspension was stored at 4°C and used within one week.

### Fabrication of miR-146a-loaded LNPs

LNPs were freshly formulated according to the above-stated protocol and used for loading pre-miR-146a (Ambion® Pre-miR™ miRNA Precursor; Thermo Fisher). Polyplexes were first generated using three different N/P ratios where *N* represents the moles of amine groups from PEI and *P* represents the moles of phosphate groups from miR-146a. The *N* values were set to 1, 10 and 20, while *P* was kept constant at 1. To make polyplexes, the appropriate amount of miR-146a was added to 50 μl of nuclease-free water in three different tubes marked as 1:1, 1:10, 1:20, followed by gentle mixing with a micropipette. Subsequently, the corresponding amounts of PEI, representing 1, 10, and 20 moles of *N*, were added to the respective tubes. The miR-146a/PEI mixtures were sonicated for 5 min at room temperature (RT) using a bath sonicator to form polyplexes. Simultaneously, 100 μl of LNPs was taken into three different tubes and sonicated for 5 min at RT to generate empty-LNPs. To generate miR-146a-loaded LNPs, the polyplex solutions were added to the corresponding LNP suspensions and further sonicated in bath sonicator for 5 min at RT. The resulting miR-146-loaded LNP formulations were kept at RT for 1 h to allow stabilization and then ultrafiltered using 50K NMWCO ultrafiltration tube (Amicon) followed by centrifugation at 5000 rpm for 15 min to remove free or unencapsulated miR-146a. The miR-146-loaded LNPs were recovered from the upper chamber of the filter tube, and the volume was adjusted to 100 μl using 20 mM HEPES buffer (pH 7.4) to obtain a final miR-146a concentration of 1 μM in 1 mg/ml LNPs. Agarose gel electrophoresis was performed to confirm miR-146a loading at different N/P ratios. For this analysis, LNP-miR146a samples were treated with 0.5% SDS for 5 min at a 1:1 ratio, mixed with 6x DNA loading buffer, and loaded onto a 2% agarose gel containing 0.2 μg/ml ethidium bromide. Gel was electrophoresed in 1x Tris-Borate-EDTA (TBE) buffer (Bio-Rad) at 100V for 90 min. To assess storage stability, samples were stored at 4°C for up to 8 days and analyzed by agarose gel electrophoresis. Based on loading efficiency and stability, the formulation prepared at an N/P ratio of 1:10 was selected for further characterization and downstream experiments.

### Physicochemical characterization of empty-LNPs and LNP-miR146a

Empty-LNPs and miR-146a-loaded LNPs were characterized to determine their hydrodynamic size and polydispersity index (PDI) in milli-Q water using dynamic light scattering (DLS), while zeta potential was measured by phase analysis light scattering (PALS) in milli-Q water (NanoBrook Omni; Brookhaven Instruments, USA). Fourier-transform infrared spectroscope (FTIR; Vertex 70, Bruker, Germany) was performed to identify functional groups in lyophilized LNP samples. The morphology of LNPs and miR-146a-loaded LNPs was assessed by transmission electron microscope (TEM; LVEM 5, Delong Instruments, Czech Republic).

### Stability study of empty-LNPs and LNP-miR146a

The stability of miR-146a encapsulated in LNPs formulated at an N/P ratio of 1:10 was investigated by agarose gel electrophoresis. Stability or degradation of miR-146a within LNPs was assessed at 37°C under different pH conditions using 20 mM HEPES buffer and in the presence of 100% mouse serum. For pH-dependent stability studies, LNP-miR146a and free miR-146a were separately mixed at a 1:1 ratio with 20 mM HEPES buffer adjusted to pH 2.5, 4.5, 7.4, 8.0, or 10.0 and incubated at 37°C for 1 h and 24 h. For serum stability study, LNP-miR146a was mixed with 100% mouse serum at a 1:1 ratio and incubated for 1 h at 37°C. Following incubation, all the treated samples were mixed with 0.5% SDS at a 2:1 ratio (two parts sample to one-part SDS) and kept at room temperature for 5 min to lyse the LNPs. Both SDS-treated and -untreated samples were then mixed with 6x DNA loading buffer and loaded onto a 2% agarose gel containing 0.2 µg/ml EtBr. Electrophoresis was performed for 90 min in 1x TBA buffer at 100V. Results were visualized and acquired using a gel documentation system.

### Animal maintenance and cell culture

C57BL/6 mice were purchased from The Jackson Laboratory (ME, USA). All animal experiments were conducted with protocols approved by the Institutional Animal Care and Use Committee (IACUC) at the University of Maryland College Park (Protocol No. R-OCT-24-36). Animals were housed in a pathogen-free environment under controlled temperature and humidity conditions, with food and water provided ad libitum. Primary thioglycolate-induced peritoneal-derived macrophages (PDMs) were isolated from 6-8 weeks old wild type (WT) C57BL/6 mice following our previously reported protocol (20). PDMs were cultured in Dulbecco’s Modified Eagle Medium (DMEM; Gibco) supplemented with 10% fetal bovine serum (FBS; Gibco) and 1x antibiotic-antimycotic solution (Thermo Fisher). Primary mouse dermal fibroblasts (MDFs) were also isolated from the C57BL/6 mice and maintained in the minimum essential medium (MEM) media, as described in our previous reports (43–46).

### Cytotoxicity testing by MTT Assay

Cytotoxicity of LNPs toward PDMs and MDFs was determined by MTT assay (47,48). Briefly, 2 x 10^5^ PDMs were seeded in 96-well plate in DMEM supplemented with 10% FBS and cultured for 24 h. The culture medium was then replaced with fresh complete DMEM supplemented with increasing concentrations of LNPs (0, 2, 10, 15, 30 and 50 μg/ml), and cells were incubated for 24 h or 48 h at 37°C. For MDF cytotoxicity study, 5 x 10^4^ cells were seeded in 96-well plates in MEM supplemented with 5% FBS and treated with the LNPs using the same concentration range and incubation times as used for PDMs. Following incubation, LNP-containing media were discarded from all the wells and replaced with serum free respective media containing 0.5 mg/ml MTT dye, followed by incubation for 3 h at 37°C to allow formation of purple formazan crystals by mitochondrial dehydrogenase enzymes. After incubation, the MTT-containing medium was discarded, and the formazan crystals were solubilized by adding DMSO. Absorbance was measured at 570 nm using a microplate reader, and percentage cell viability was calculated as previously described.

### LIVE/DEAD viability assay

PDMs (4 x 10^5^ cells) and MDFs (1 x 10^5^ cells) were seeded onto 12 mm sterile glass coverslips in 24-well plates containing their respective culture media, as mentioned above, and incubated for 48 h at 37°C (46). Following incubation, the medium was replaced with complete medium supplemented with 50 μg/ml LNPs, and cells were further incubated for 48 h or 96 h at 37°C. Untreated wells served as controls. After incubation, wells were washed twice with 1 x PBS and stained according to the manufacturer’s instruction using the LIVE/DEAD® Viability/Cytotoxicity Assay Kit (Invitrogen; L3224). Fluorescence microscopy images were obtained and processed using Fiji ImageJ software to quantify cell viability.

### Cellular uptake of NBD-labeled LNPs

To investigate cellular uptake of LNPs in PDMs and MDFs, NBD-PE-tagged LNPs were formulated by replacing 2 mM DOPE with 2 mM NBD-PE, following the above-stated LNP formulation method. The molar ratio of the lipid components (DOPE:DOTAP:LA:TPGS:NBD-PE) was 38:10:48:2:2, respectively. The lipid mixtures were loaded into a Hamilton syringe and injected into 20 mM HEPES buffer under continuous stirring, followed by incubation in the dark for 1 h. PDMs (4 x 10^5^ cells) and MDFs (1 x 10^5^ cells) were seeded onto 12 mm sterile glass coverslips in 24-well plates containing their respective culture media and incubated for 48 h at 37°C. Following incubation, the medium was discarded, and fresh complete medium containing 50 μg/ml NBD-PE-tagged LNPs (LNP-NBD-PE) was added, and cells were further incubated for 1 h at 37°C. Following treatment, cells were washed twice with ice-cold 1 x PBS to halt further internalization of LNP-NBD-PE. Cellular uptake of LNP-NBD-PE was visualized using a fluorescence microscope (Zeiss Axio Observer Z1 Motorized Inverted Fluorescence Microscope). Acquired images were processed using Fiji ImageJ software.

### Uptake of Cy3-labeled scrambled-miR146a-loaded LNPs

To assess the transfection efficiency and intracellular stability of LNP-miR146a over time, Cy3-labeled scrambled-miR146a was loaded into LNPs at an N/P ratio of 10 using the protocol described above and used immediately. PDMs (4 x 10^5^ cells) and MDFs (1 x 10^5^ cells) were seeded onto 12 mm sterile glass coverslips in 24-well plates containing their respective culture media and incubated for 48 h at 37°C. Following incubation, the culture medium was replaced with fresh complete medium, and cells were transfected with LNP-Cy3-Scr-miR146a at a final concentration of 50 nM. Cells were incubated for 1, 2 and 24 h at 37°C. After incubation, the medium was removed, and cells were washed twice with 1 x PBS and fixed with 4% paraformaldehyde, followed by additional washes with 1 x PBS. Coverslips were mounted using a DAPI-containing solution and allowed to cure overnight at room temperature in the dark. The presence of LNP-Cy3-Scr-miR146a and nuclei was visualized using a fluorescence microscope. Images were captured and processed using Fiji ImageJ software.

### Endo/lysosomal escape study

PDMs (4 x 10^5^ cells) and MDFs (1 x 10^5^ cells) were seeded onto 12 mm glass coverslips in 24-well plates and incubated at 37°C for 48 h. The culture medium was then replaced with fresh complete medium containing 50 nM LysoTracker™ Green DND-26 (Invitrogen) to stain lysosomes. Cells were subsequently treated with 50 nM LNP-Cy3-Scr-miR146a and incubated for 2 h at 37°C. For the 24 h time-point study, cells were first treated with 50 nM LNP-Cy3-Scr-miR146a, and at 22 h post-treatment, 50 nM LysoTracker was added and incubated for an additional 2 h at 37°C. After incubation, the medium was discarded, and cells were washed twice with 1 x PBS, fixed with 4 % paraformaldehyde, and washed again with 1 x PBS. Glass coverslips were carefully removed and inverted on rectangular glass slides containing a DAPI-containing mounting medium, then kept in the dark overnight to allow solidification. Images were acquired using a confocal microscope (FLUOVIEW FV3000) with DAPI (blue), FITC (green), and TRITC (red) filter channels to visualize nuclei, lysosomes, and Cy3-labeled LNPs, respectively. Image analysis was performed using Fiji ImageJ software.

### In vivo cytotoxicity analysis

To investigate in vivo biocompatibility, 6-8 weeks old WT C57BL/6 mice were selected and divided in three groups (n = 3 per group): HEPES, LNPs, and LNP-miR146a. LNPs were injected intravenously via the tail vein at a dose of 1 mg/kg body weight. The miR-146a concentration was 50 nM per 50 μg of LNPs. Control mice received 50 μl of HEPES buffer only. All the mice were housed in a sterile environment for 7 days. After the incubation period, mice were euthanized, and blood was collected from heart for further analysis. Blood was divided in two parts to obtain serum and whole blood. Serum was used to assess hepatic function markers - aspartate aminotransferase (AST) and alanine transaminase (ALT); kidney function markers - blood urea nitrogen (BUN) and creatinine; and basic metabolic electrolytes - calcium, bicarbonate, chloride, potassium, and sodium. Whole blood was used to assess hematological parameters including white blood cells (WBC) count, platelet count, hemoglobin, and hematocrit. Collected samples were sent to IDEXX BioAnalytics (North Grafton, MA, USA) for analysis. The resulting data were analyzed and plotted using GraphPad Prism.

### Analysis of subcutaneous retention and systemic biodistribution of LNPs in vivo

In vivo subcutaneous (s.c) retention and organ distribution were investigated in WT C57BL/6 mice using an In Vivo Imaging System (IVIS). Cy5.5-PE (Cyanine 5.5 PE; 810336; Avanti Polar lipids, USA)-labeled LNPs were formulated following the above-mentioned LNP formulation method. The molar ratio of lipid components (DOPE:DOTAP:LA:TPGS) was 40:10:48:2, respectively. Cy5.5-PE (1 μg/µl) was mixed with the lipid mixture, loaded into a Hamilton syringe, and injected into 20 mM HEPES buffer under stirring, followed by incubation in the dark for 1 h. The final Cy5.5 concentration was adjusted to 10 μg/ml, which provided optimal excitation and emission as per manufacturer’s specifications. To visualize the distributions of Cy5.5-LNPs, mice were anesthetized, and the dorsal skin was shaved prior to imaging. Cy5.5-LNPs were injected s.c into the dorsal skin at the volume of 50 μl (10 μg/ml). IVIS images were taken immediately after injection (day 0) and subsequently on days 3, 7, and 14 (n = 3 per group) using an IVIS Spectrum system (PerkinElmer, USA). On day 14, mice were euthanized, and organs including dorsal skin, dorsal muscles, liver, spleen, lungs, heart, and kidneys were harvested to assess the biodistribution and migration of Cy5.5-LNPs using IVIS imaging and analyzed by LivingImage software.

### Analysis of in vivo delivery and uptake of Cy5.5-LNP, LNP-Cy3-Scr-miR146a, and HEPES/Cy3-Scr-miR146a to skin tissue and other organs in mice

Eight-week-old ApoE^-/-^(C57BL/6 background) mice were purchased from The Jackson Laboratory and housed as per IACUC guideline in the UMD animal facility. Mice were maintained on a high-fat diet for 6 weeks, after which the uptake and biodistribution of Cy5.5-LNP nanoparticles were evaluated. Cy5.5-LNP (50 μl containing 10 μg/ml Cy5.5) were intravenously injected (n = 3 per group), and mice were euthanized 3 h post-injection, and organs were harvested for analysis. The heart with the aortic arch and the full abdominal aorta were carefully collected and placed in sterile petri dishes. Additional organs, including lungs, liver, kidneys, and spleen, were also collected and placed in separate petri dishes. Uptake and biodistribution of Cy5.5-LNPs in all harvested organs were visualized using an IVIS and analyzed by LivingImage software.

Skin tissue uptake, distribution, and stability of Cy3-tagged scrambled-miR146a loaded in LNPs, as well as control HEPES/Cy3-Scr-miR146a (dye-labeled miR-146a alone), were evaluated in 6-8-weeks-old WT mice using IVIS. Mice were divided into two groups (n = 3 per group), and the dorsal skin was shaved to minimize autofluorescence interference. One group received s.c injection of LNP-Cy3-Scr-miR146a (50 nM), while the control group received HEPES/Cy3-Scr-miR146a (50 nM). IVIS images were collected at 0 h, and 3 h post-injection, and subsequently on days 1, 4, and 7. An additional cohort of mice (n = 3 per group) was euthanized on day 4 post-injection, and organs (skin, liver, lungs, heart, spleen, and kidneys) were harvested, placed in sterile petri dishes, and imaged using an IVIS. All images were analyzed using LivingImage software.

### Quantitative real-time RT-PCR (RT-qPCR)

Bone marrow-derived macrophages (BMDMs) were prepared and cultured according to our previously published protocol (20). Approximately 2 x 10^6^ BMDMs were seeded in 60-mm culture dishes in DMEM supplemented with 10% FBS and incubated for 24 h at 37°C. After cell adhesion, the medium was replaced with fresh complete DMEM. miR-146a and scrambled-miR-146a oligonucleotides were loaded into LNPs at an N/P ratio of 10 prior to use. BMDMs were treated with 50 nM LNP-miR146a or LNP-Scr-miR146a in complete DMEM and incubated for 24 h at 37°C. Following treatment, the medium was discarded, cells were washed twice with 1 x PBS, and cells were harvested form each dish. Total RNA and microRNA were extracted from BMDMs according to the manufacturer’s instructions (Qiagen), followed by treatment with RNase-free DNase. The concentration and purity of the extracted RNA and microRNA were determined using a Nanodrop-2000 system (Thermo Fisher Scientific).

To evaluate miR-146a expression, cDNA was synthesized using the miRCURY LNA SYBR Green PCR Kit according to the manufacturer’s instructions (Qiagen) using a SimpliAmp Thermal Cycler (Applied Biosystem). RT-qPCR was then performed using the miRCURY LNA SYBR Green PCR Kit with primers for hsa-miR-146a-5p (YP00204688) and SNORD68 (YP00203911), following the manufacturer’s instructions (Qiagen). Amplification was carried out for 40 cycles using a CFX96 cycler (Bio-Rad). Threshold cycle (Ct) values for hsa-miR-146a-5p were normalized to the average Ct value of SNORD68, and relative miR-146a expression was calculated using the ΔΔCt method. In addition, expression of TRAF6 was assessed in LNP-Scr-miR146a- and LNP-miR146a-treated BMDMs. cDNA synthesized using iScript Advance cDNA Kit (172–5037) and RT-qPCR was performed using the SsoAdvanced Universal SYBR Green Supermix (172–5270) and primers for TRAF6 (qMmuCEP0054569) and GAPDH (qMmuCID0018612), according to the manufacturer’s instructions (Bio-Rad). Reactions were run for 40 cycles on a CFX96 cycler (Bio-Rad). Ct values of target genes were normalized to GAPDH, and relative gene expression was calculated using the ΔΔCt method.

### In vivo LNP-mediated miR-146a overexpression and inflammatory gene expression analysis in a biomaterial implantation model

To assess functional overexpression of miR-146a in biomaterial-implanted skin tissue, 6-8-week-old WT mice were divided into two groups (n = 4 per group) and injected with either LNP-miR146a or LNP-Scr-miR146a. Cellulose ester membranes (0.22 μm pore size; Millipore-Sigma) were cut into 10-mm-diameter discs and sterilized by autoclaving. Sterilized membranes were submerged in 100% mouse serum and incubated overnight at 4°C for surface coating. For membrane implantation, dorsal hair was shaved, and a small incision was created using a sterile scalpel and scissors, followed by insertion of the implants between dorsal skin and underlying muscle. The incision was closed using absorbable suture. After surgery, 50 nM LNP-miR146a or LNP-Scr-miR146a was injected locally near the implanted site in the respective groups on days 0 and 14. After 28 days of implantation, mice were euthanized and skin tissue containing the implant was harvested for analysis. Homogenized tissues were processed to harvest both microRNA and total RNA according to the manufacturer’s instructions (Qiagen), followed by treatment with RNase-Free DNase. RNA concentration and purity were assessed using a Nanodrop-2000 system. Expression levels of miR-146a, TRAF6, IL-6 (qMmuCED0045760), and TNF-α (qMmuCED0004141) were evaluated and analyzed following the above-stated protocol.

### Foreign body giant cell formation

Foreign body giant cell (FBGC) formation was evaluated using BMDMs following our previously published protocol (20). Briefly, 3 x 10^5^ BMDMs were seeded onto 8-well Permanox plastic chamber slides in 10% FBS-containing DMEM, and incubation at 37°C for 48 h. After cell adhesion, old medium was replaced with fresh complete DMEM, and cells were treated with LNP-miR146a or LNP-Scr-miR146a (50 nM) for 24 h. Following the 24 h treatment, the medium was supplemented with IL-4 (25 ng/ml) and GMCSF (25 ng/ml), and replaced every 48 h, and cells were maintained under the same conditions for an additional 96 h. BMDMs treated with only IL-4 and GMCSF alone served as controls. After 96 h of IL-4 and GMCSF treatment, cells were washed twice with 1 x PBS and fixed with 4% paraformaldehyde for 10 min, followed by additional washes with 1 x PBS. Fixed cells were stained with 1 x Giemsa stain to visualize cell nuclei. Images were acquired using a Nikon fluorescence microscope (Nikon Eclipse Upright) and analyzed using Fiji ImageJ software.

### Declaration of generative AI and AI-assisted technologies in the writing process

During the preparation of this work the author(s) used ChatGPT 5.2 in order to improve readability and perform proofreading. After using this tool/service, the author(s) reviewed and edited the content as needed and take(s) full responsibility for the content of the publication.

## Results

### Formulation and physicochemical characterization of empty-LNPs and LNP-miR146a

LNPs were formulated by injecting an ethanolic lipid phase into an aqueous buffer under constant stirring. The rapid transition of the aqueous solution into an opalescent dispersion indicated spontaneous self-assembly of LNPs, as shown in the Figure 1A. The functional roles of individual lipids involved in LNP assembly are schematically shown in Figure 1B. Samples were withdrawn at 15, 30, and 60 min to determine changes in the hydrodynamic diameter of the LNPs. The particle sizes were approximately 166, 159 and 144 nm, respectively, whereas the size distribution of 60-min filtered sample was 145 nm (Figure 1C). Moreover, the PDI improved with increased stirring time and subsequent filtration. The PDI values of 15-min and 30-min stirred samples were 0.026 and 0.076, respectively, while the 60-min stirred sample showed a PDI of 0.119, which further improved to 0.126 following filtration. These results indicate that size uniformity and PDI of the LNPs improved with prolonged stirring followed by filtration. Furthermore, the surface charge of the LNPs was assessed using Zeta potential measurements via PALS and was found to be approximately −24 ± 3 mV (Figure 1D). FTIR spectroscopy confirmed the presence of characteristic lipid functional groups, including C-H stretching, C=O (ester carbonyl), and C-O vibrations, verifying the chemical integrity of the lipid components and successful nanoparticle assembly (Figure 1E). In addition, changes in the physicochemical properties of LNP-miR146a were observed compared to empty-LNPs, with the results summarized in Figure 1F. The size distribution, PDI, zeta potential, and encapsulation efficiency of LNP-miR146a were 182.93 ± 3.38 nm, 0.168 ± 0.024, −6.21 ± 1.25 mV, and 95 ± 2.80%, respectively. In contrast, the size distribution, PDI, and zeta potential for empty-LNPs were 163.79 ± 7.69 nm, 0.122 ± 0.050, and - 21.42 ± 4.51 mV, respectively. Transmission electron microscopy (TEM) images (Figure 1G) confirmed the spherical morphology and nanoscale structure of the LNPs and indicates that particle morphology of LNPs remained unchanged following miR146a loading. Enlarged views (highlighted by red boxes) reveal uniform, smooth-surfaced particles, consistent with the DLS size measurements.

**Figure 1.**
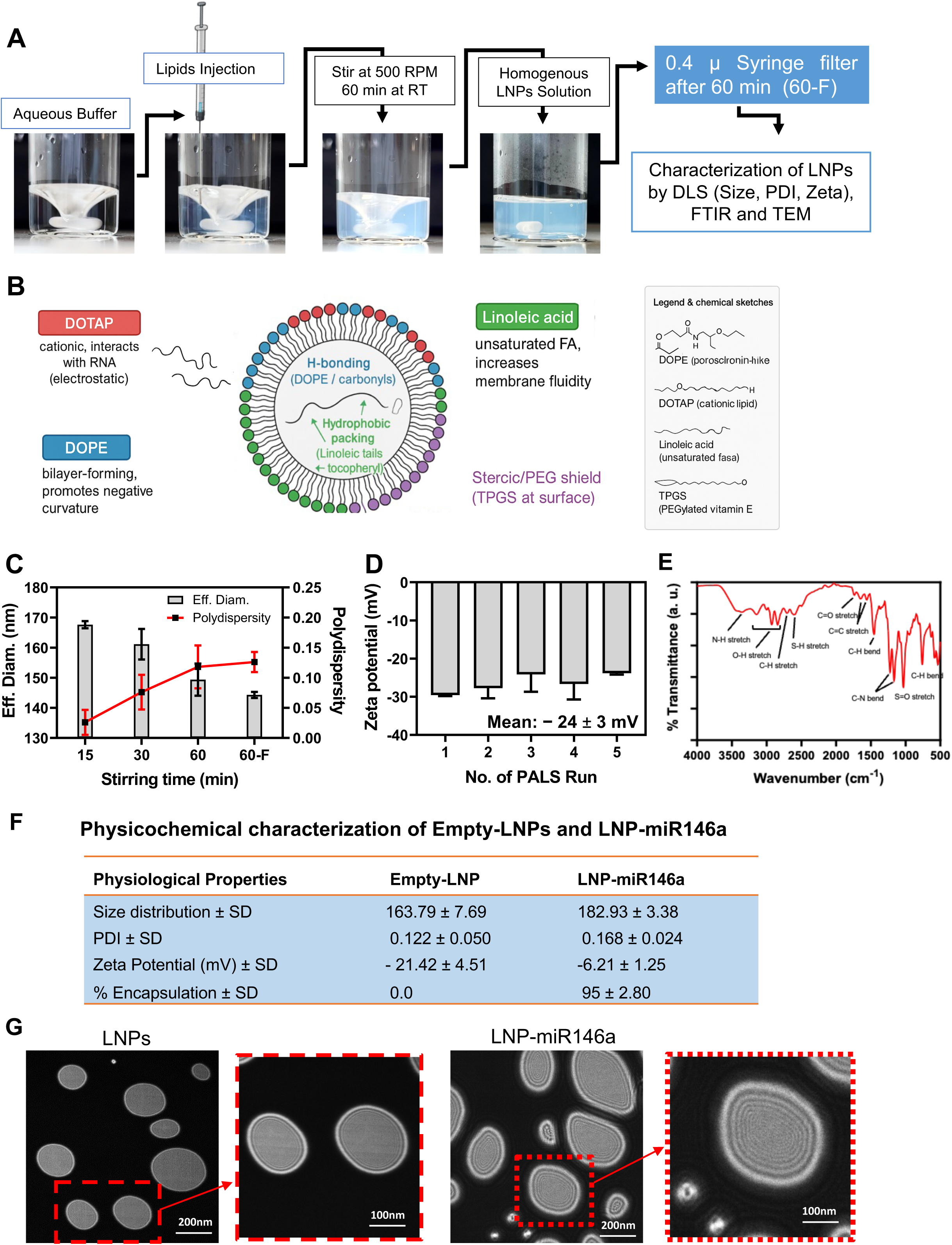
Engineering and physicochemical characterization of empty-LNPs and miR-146a-loaded LNPs. **(A)** LNPs were formulated using an ethanolic lipid injection method into an aqueous buffer followed by stirring. **(B)** Schematic illustration of the assembly of lipid components used to prepare the LNPs. The characteristic properties of the LNPs were determined using multiple analytical techniques, including **(C)** DLS, **(D)** PALS and **(E)** FTIR. The table summarizes the physicochemical properties of empty-LNPs and LNP-miR146a, including particles size, PDI, zeta potential, and encapsulation efficiency. **(G)** The morphology of empty-LNPs and LNP-miR146a was evaluated by transmission electron microscopy. All formulations were prepared in triplicate, and the data are presented as the mean ± standard error of the mean.

miR-146a loading into LNPs was performed using three different N/P ratios, and entrapment of miR-146a was confirmed by agarose gel electrophoresis. A schematic illustration of miR-146a loading into LNPs and its separation on agarose gel is shown in Figure 2A. miR-146a-loaded LNPs prepared at different N/P ratios (1:1, 1:10, and 1:20) were subjected to ultrafiltration, and the collected filtrates were analyzed by agarose gel to check the presence of unencapsulated miR-146a (Figure 2B). Among the tested formulations, the N/P ratio of 1:10 showed the highest entrapment efficiency. LNP-miR146a formulations were further treated with 0.5% SDS at a 1:1 ratio and analyzed by agarose gel electrophoresis. The resulting bands confirmed successful miR-146a loading and its retention within the LNPs (Figure 2C), as well as stability during storage at 4°C for up to 8 days post-formulation (Figure 2D).

**Figure 2.**
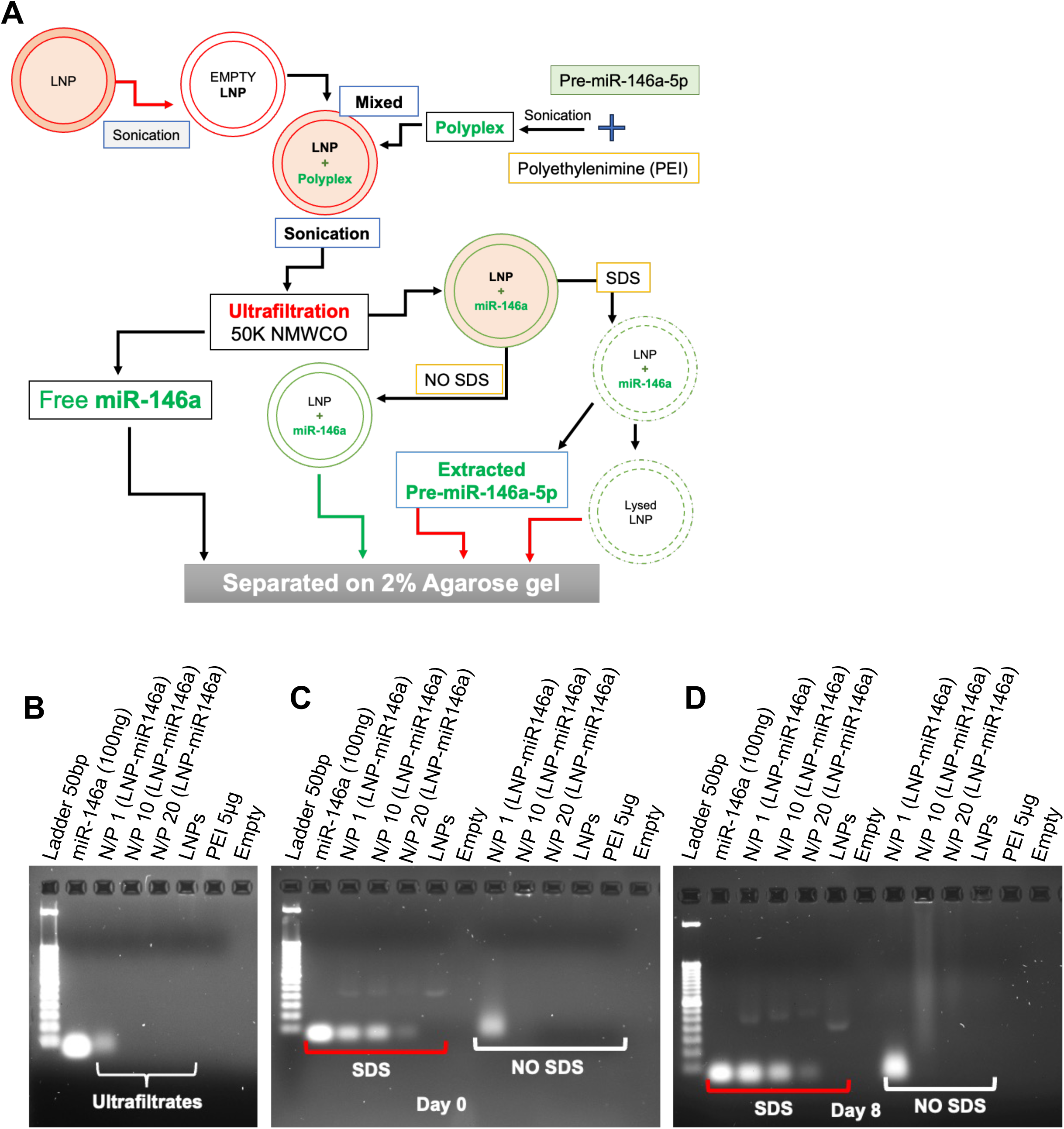
miR-146a Loading and Encapsulation Efficiency in Lipid Nanoparticles. **(A)** Schematic representation of miR-146a loading into LNPs and its validation by agarose gel electrophoresis. LNP-miR146a was formulated using three different N/P ratios. **(B)** Ultrafiltrates were separated by agarose gel electrophoresis to assess unencapsulated miR-146a. **(C)** LNP-miR146a treated with SDS on day 0 was analyzed by agarose gel electrophoresis to determine the optimum N/P ratio. **(D)** The stability of miR146a was determined by storing the formulations at 4°C for 8 days, followed by agarose gel electrophoresis.

The stability of miR-146a encapsulated in LNPs was further investigated under varying pH conditions at 37°C and analyzed by agarose gel electrophoresis. As shown in Figure 3, LNP disruption and miR-146a release were observed at high pH (pH 10.0), evidenced by the appearance of miR-146a bands in SDS-untreated samples after 1 h and 2 h of incubation at 37°C. After 1 h of incubation at 37°C, miR-146a bands were detected at all pH values following 0.5% SDS treatment, indicating that LNP-miR146a remained stable under both acidic and mildly basic conditions (Figure 3A). However, a reduction in band intensity was observed at pH 2.5 and 4.5 after 24 h of incubation at 37°C, suggesting partial degradation under strongly acidic conditions (Figure 3B).

**Figure 3.**
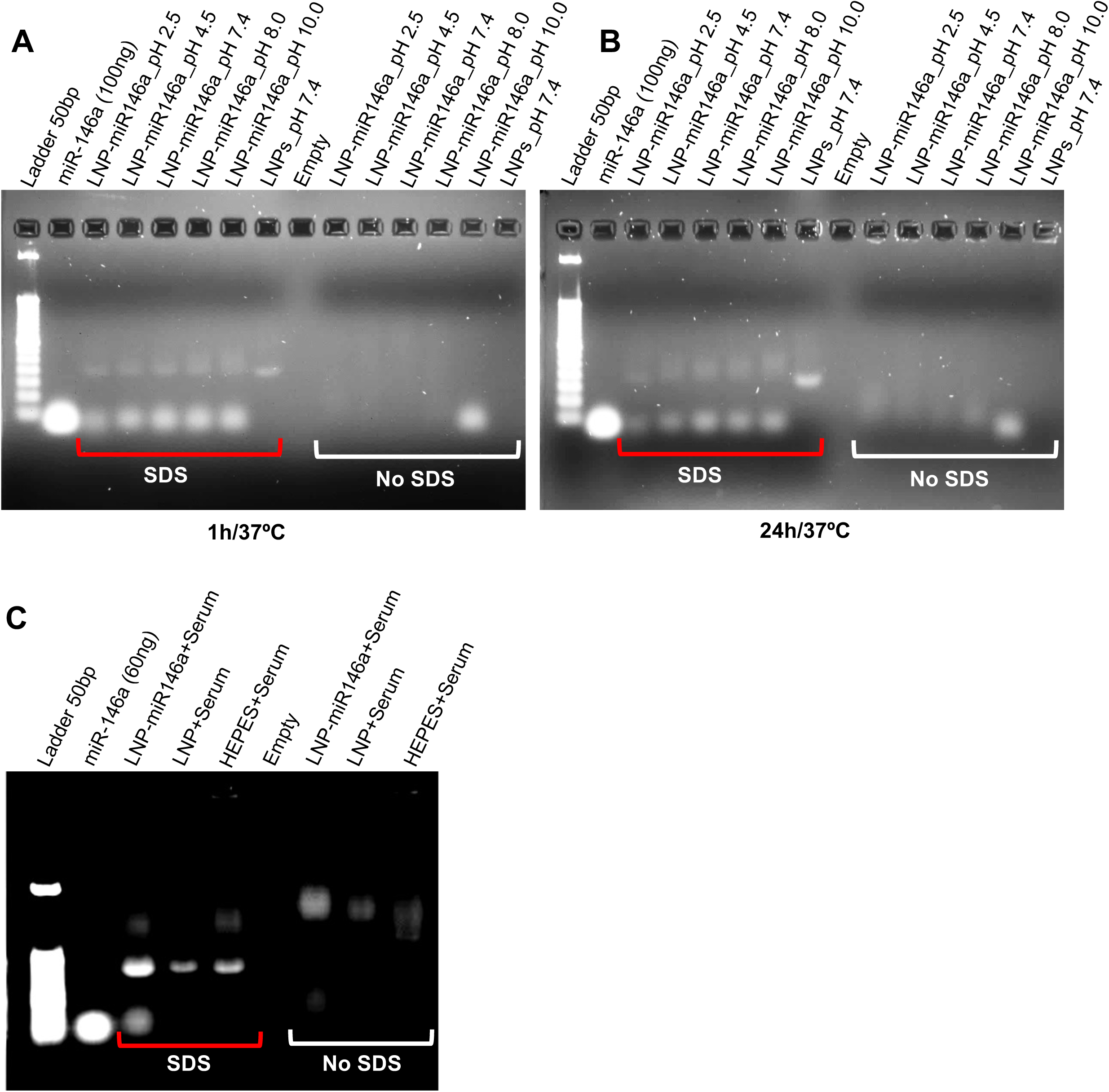
Stability of LNP-Encapsulated miR-146a under pH and Serum Conditions. The stability of miR-146a encapsulated in LNPs was checked under different pH conditions after **(A)** 1 h and **(B)** 24 h of incubation at 37°C. **(C)** In addition, the stability of miR-146a in LNPs was evaluated in the presence of 50% serum at 37°C and analyzed by agarose gel electrophoresis after 1 h of incubation.

In addition, miR-146a stability in LNPs was checked by incubating the formulation with 100% serum at a 1:1 ratio, followed by agarose gel analysis after 1 h at 37°C. miR-146a bands were detected only in SDS-treated samples, while no bands were observed in SDS-untreated samples, confirming protection of miR-146a within LNPs in the presence of crude mouse serum (Figure 3C).

Overall, these results demonstrate the successful preparation of stable, uniformly sized empty-LNPs and LNP-miR146a using a simple lipid injection and stirring method. Filtration serves as a critical refinement step that enhances physicochemical stability and homogeneity of this LNPs formulation, supporting the suitability of this LNP platform for downstream biomedical or drug-delivery applications.

### Excellent biocompatibility of LNPs in both PDMs and MDFs

Assessing the biocompatibility of LNPs is critical for their biomedical application. Accordingly, we checked LNP cytocompatibility in two normal murine primary cell types - PDMs and MDFs - by assessing cell viability. The LNPs exhibited favorable biocompatibility toward both PDMs and MDFs after 24 and 48 h of exposure across increasing concentrations. Approximately 80% viability of PDMs was detected at 24 h in the presence of 50 μg/ml LNPs (Figure 4A), and a comparable level of viability was maintained after 48 h of incubation at the same concentration (Figure 4B). LNPs showed even greater biocompatibility toward MDFs, with nearly 100% cell viability observed at all tested concentrations after 24h of treatment (Figure 4C). After 48 h of incubation, a slight reduction in MDF viability was observed at 30 and 50 μg/ml, however, viability remained close to 90% (Figure 4D). To further validate these findings, live/dead assays were performed in both PDMs and MDFs following 48 and 96 h of incubation with 50 μg/ml LNPs. As shown in Figure 4E-F, very few dead (red) PDMs were found relative to live (green) cells in LNP-treated samples. Similarly, LNPs were well tolerated by MDFs, and no detectable dead cells observed after 48 and 96 h of treatment (Figure 4G-H). Collectively, these data confirm the excellent biocompatibility of LNPs in both PDMs and MDFs. Based on these results, a concentration of 50 μg/ml LNPs was selected for subsequent in vitro studies.

**Figure 4.**
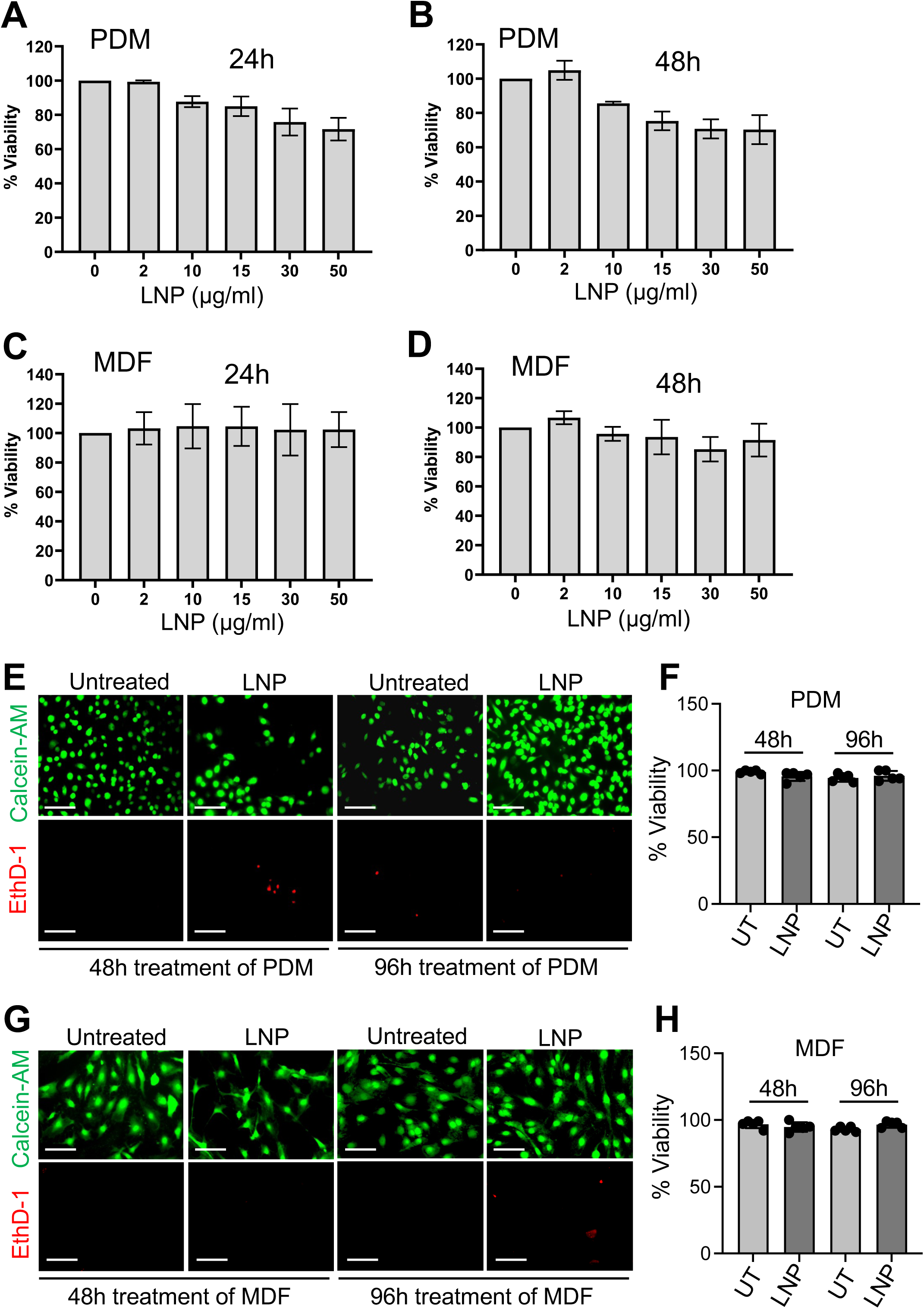
In vitro biocompatibility assessment of LNPs. Cytotoxicity of LNPs was examined using an MTT assay in PDMs after 24 h **(A)** and 48 h **(B)** of incubation with increasing concentrations of LNPs. Similarly, cytotoxicity of LNPs was checked in MDFs after 24 h **(C)** and 48 h **(D)** of treatment. Cell viability of PDMs **(E, F)** and MDFs (**G, H**) was further checked using a LIVE/DEAD assay by fluorescence microscope after 48 h and 96 h of treatment with 50 µg/ml LNPs. All experiments were performed in triplicate, and data are represented as the mean ± standard error of the mean.

### Efficient intracellular uptake of empty-LNPs and LNP-miR146a in PDMs and MDFs

The cellular uptake of NBD-PE-labeled LNPs was evaluated in PDMs and MDFs by fluorescence microscopy to assess nanoparticle internalization (Figure 5A). Uptake of LNP-NBD-PE was significantly higher in PDMs than MDFs (Figure 5B), consistent with the phagocytic nature of macrophages. In addition, the uptake kinetics of Cy3-labeled scrambled miR-146a-loaded LNPs (LNP-Cy3-Scr-miR146a) were quantified in both PDMs and MDFs in a time-dependent manner (1, 2, and 24 h). In PDMs, comparable levels of intracellular fluorescence were observed across all time points, indicating rapid internalization, with near-complete uptake occurring within 1 h and remaining stable up to 24 h (Figure 5C). No substantial reduction in Cy3 fluorescence intensity was detected at 24 h of incubation at 37°C, suggesting intracellular stability of LNP-Cy3-Scr-miR146a (Figure 5D). In contrast, uptake of LNP-Cy3-Scr-miR146a in MDFs occurred more gradually, with complete internalization observed after 24 h of incubation (Figure 5E-F).

**Figure 5.**
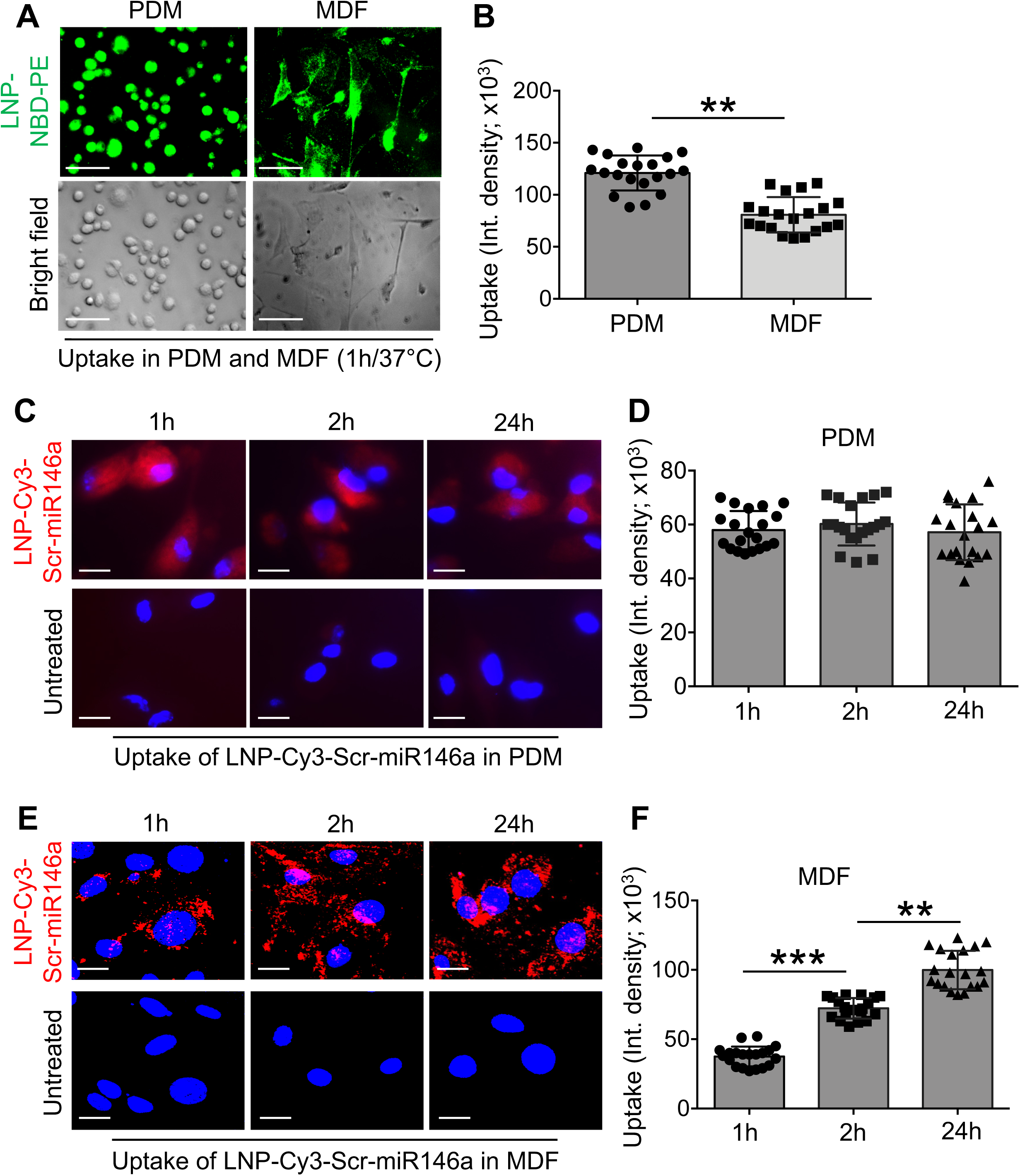
Uptake of empty-LNPs and LNP-miR146a in PDMs and MDFs. The uptake of LNPs was assessed in PDMs and MDFs using NBD-PE-tagged LNPs by measuring NBD fluorescence intensity in both cells **(A, B)**. Uptake of LNP-Cy3-Scr-miR146a was evaluated in PDMs **(C, D)** and MDFs **(E, F)** after 1, 2, and 24 h of incubation at 37°C. All experiments were performed in triplicates; represented images are shown, quantitative data are presented as the mean ± standard error of the mean (SEM). Statistical significance was determined using one-way ANOVA and is denoted as **p<0.01, ***p<0.001.

Altogether, these results suggest that both empty-LNPs and miR-146a-loaded LNPs are efficiently internalized by both non-phagocytic and phagocytic cells, highlighting their potential utility as versatile carriers for intracellular miR delivery.

### Endo/lysosomal escape of LNP-miR146a in PDMs and MDFs

Efficient endo/lysosomal escape is essential for LNP-mediated delivery of genetic cargo, as it prevents lysosomal degradation and enables cytosolic release of nucleic acids. To evaluate the endo/lysosomal escape capability of the formulation, LNP-Cy3-Scr-miR146a was internalized into PDMs and MDFs, and intracellular trafficking was analyzed by confocal fluorescence microscopy (Figure 6). In PDMs, substantial colocalization of LNP-Cy3-Scr-miR146a (red) with lysosomes (green) was observed at 2 h post-incubation. This colocalization markedly decreased by 24 h, as evidenced by the spatial separation of red and green fluorescence signals (Figure 6A). Quantitative analysis of colocalization (yellow signal) revealed an approximately 70% reduction at 24 h compared with 2 h, indicating efficient lysosomal escape over time (Figure 6B). A similar trend was observed in MDFs. At 2 h, LNP-Cy3-Scr-miR146a showed higher lysosomal colocalization, although lysosomal staining appeared less distinctly resolved. By 24 h, clear separation of lysosomal (green) and nanoparticle (red) signals was detected, consistent with successful endosomal/lysosomal escape (Figure 6C). Quantitative analysis demonstrated an approximately 80% reduction in colocalization intensity at 24 h relative to 2 h (Figure 6D). Collectively, these results demonstrate that LNP-miR146a exhibits robust endo/lysosomal escape in both phagocytic and non-phagocytic cells, underscoring its potential as an effective intracellular delivery platform for therapeutic nucleic acids and drugs.

**Figure 6.**
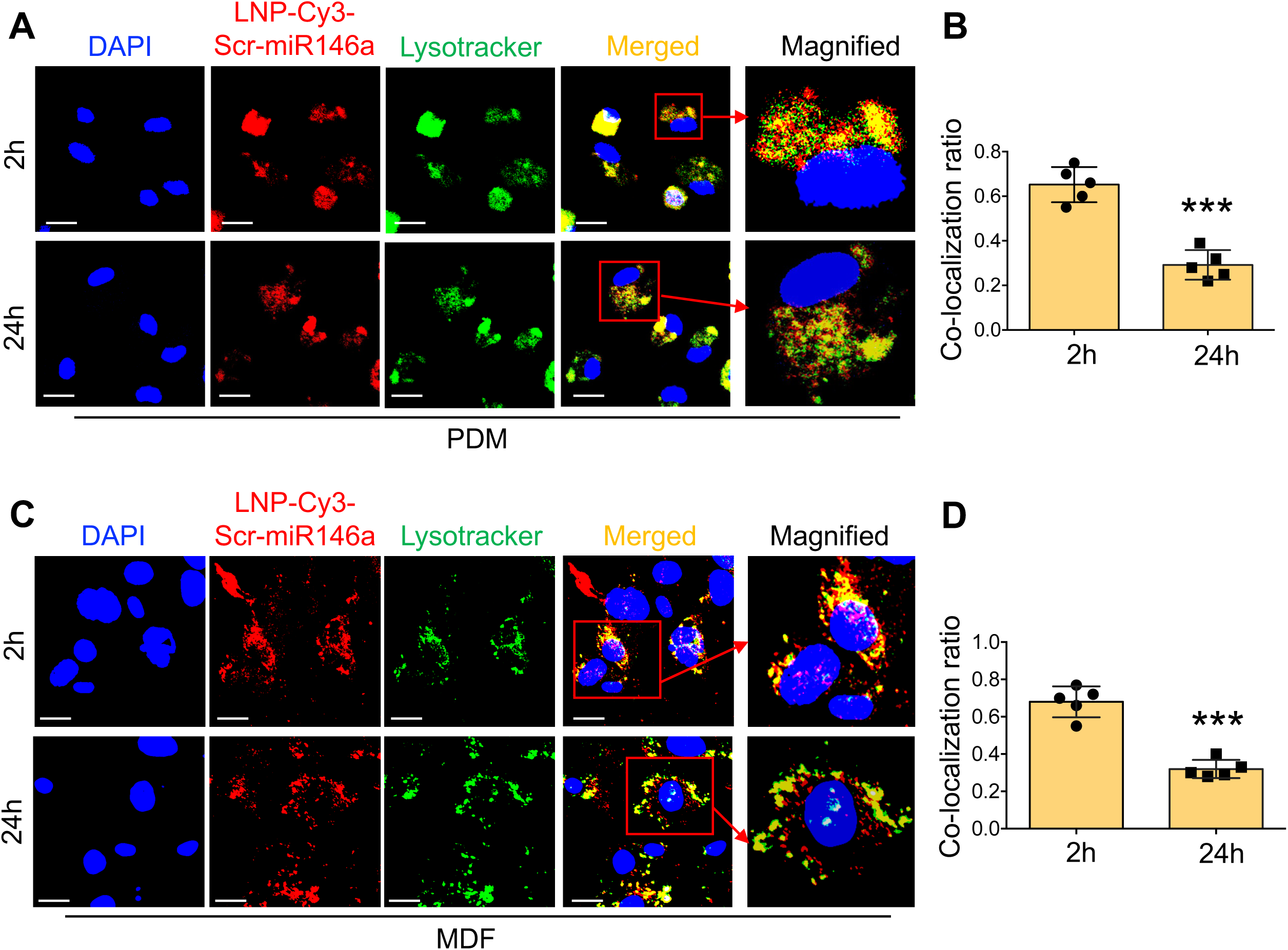
Endo/Lysosomal escape of miR-146a-loaded LNPs. Endo/lysosomal escape of LNP-Cy3-Scr-miR146a was evaluated in PDMs after 2 h and 24 h of treatment **(A).** Escape efficiency was quantified by calculating the colocalization ratio based on the overlap of red (LNP-Cy3-Scr-miR146a) and green (lysosomal) fluorescence signals, which appear yellow in merged images **(B).** A similar analysis was performed in MDFs **(C),** and colocalization ratios were quantified at 2 h and 24 h of incubation based on yellow fluorescence intensity **(D)**. All experiments were performed in triplicate; representative images are shown, and quantitative data are presented as the mean ± standard error of the mean. Statistical significance was determined using ANOVA and is denoted as ***p < 0.001.

### Empty-LNPs and LNP-miR146a exhibits no detectable toxicity in vivo

The in vivo cytotoxicity of empty LNPs and LNP-miR146a was evaluated to assess their systemic biocompatibility. Hepatic function was examined by measuring serum AST and ALT levels. AST levels were approximately 200 U/L in both HEPES- and LNP-treated mice, while a modest increase was observed in the LNP-miR146a-treated group; however, values remained within the acceptable range of 200-250 U/L (Figure 7A). ALT levels across all treatment groups ranged from 30 to 150 U/L, which is within the normal physiological range for healthy WT mice (Figure 7B). These results indicate normal hepatic function following LNP administration. Renal function was assessed by measuring BUN and creatinine levels. BUN values for all treated groups were within the range of 20-30 mg/dL, consistent with healthy renal function and indicative of no detectable kidney toxicity (Figure 7C). Similarly, serum creatinine levels were maintained between 0.10 and 0.15 mg/dL across all treatment groups, further confirming normal renal function (Figure 7D). Metabolic electrolyte levels, including sodium, potassium, bicarbonate, and chloride, were also evaluated. No abnormal elevations or reductions were observed in any treatment group, and all electrolyte concentrations remained within the normal physiological range for healthy mice (Figures 7E-I).

**Figure 7.**
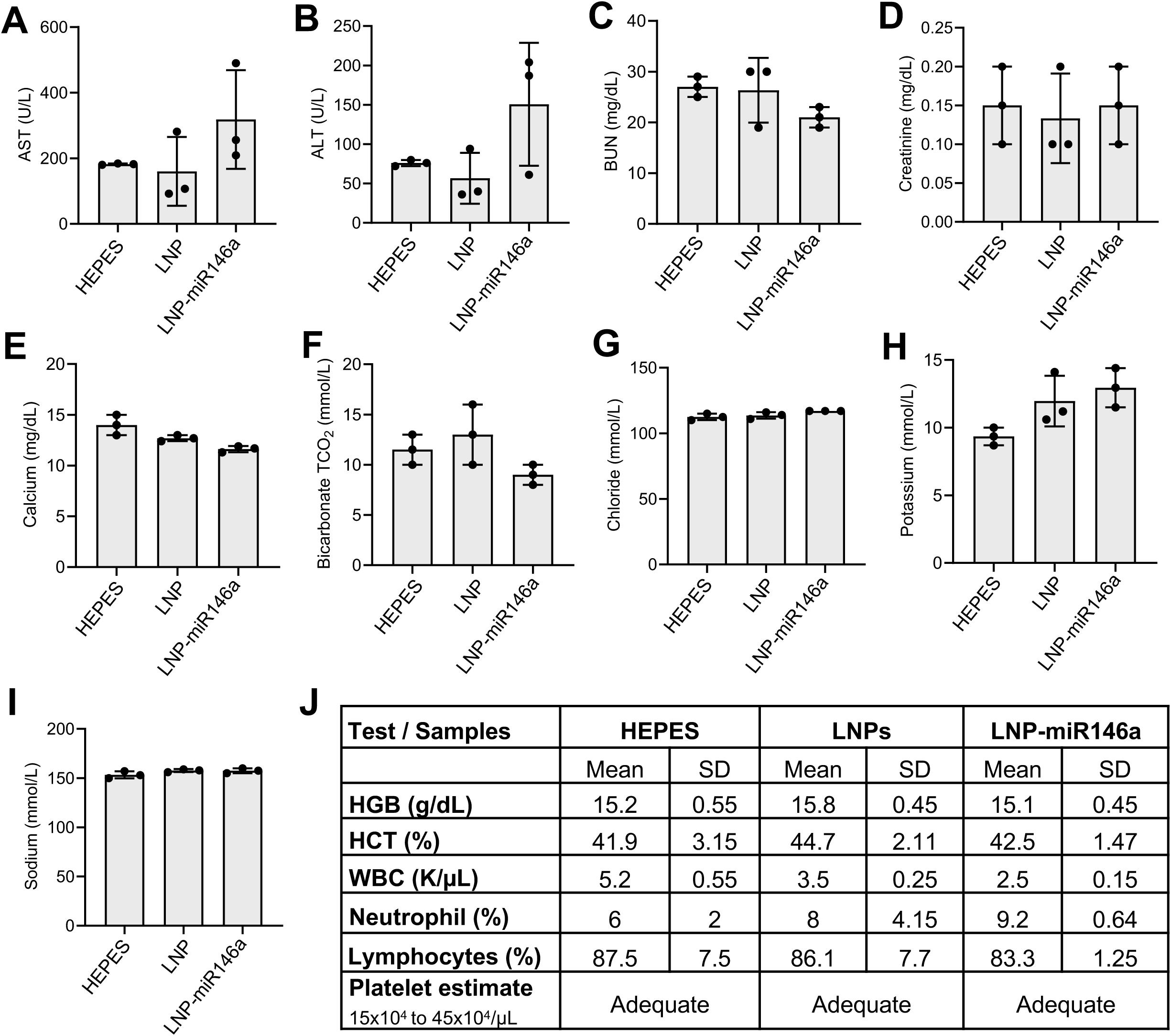
Systemic biocompatibility of miR-146-loaded LNPs in vivo. In vivo cytotoxicity was evaluated in mice by examining hepatic function markers, including AST **(A)**, and ALT **(B)**, as well as kidney function markers, including BUN **(C)** and creatinine **(D)**. Basic metabolic electrolytes - calcium **(E)**, bicarbonate **(F)**, chloride **(G)**, potassium **(H),** and sodium **(I) –** were also measured. **(J)** Hematological parameters were analyzed in whole blood samples. All experiments were performed in triplicates, and data are represented as the mean ± standard error of the mean.

Hematological parameters were analyzed using whole blood samples and were found to be comparable across all groups (Figure 7J). In HEPES-treated mice, hemoglobin (HGB), hematocrit (HCT), WBC count, neutrophil percentage, lymphocyte percentage, and platelet levels were 15.2 ± 0.55 g/dl, 41.9 ± 3.15%, 5.2 ± 0.55 K/μl, 6 ± 2%, 87.5 ± 7.5%, and adequate, respectively. Corresponding values in LNP-treated mice were 15.8 ± 0.45 g/dl, 44.7 ± 2.11%, 3.5 ± 0.25 K/μl, 8 ± 4.14%, 86.1 ± 7.7%, and adequate platelet counts. Similarly, LNP-miR146a-treated mice exhibited HGB, HCT, WBC, neutrophil, lymphocyte, and platelet values of 15.1 ± 0.45 g/dl, 42.5 ± 1.47%, 2.5 ± 0.15 K/μl, 9.2 ± 0.64%, 83.3 ± 1.25%, and adequate platelet levels, respectively.

Collectively, these findings demonstrate that both empty LNPs and LNP-miR146a nanoparticles exhibit excellent in vivo biocompatibility, with no evidence of hepatic, renal, metabolic, or hematological toxicity, supporting their suitability for therapeutic applications.

### In vivo delivery of empty-LNPs and LNP-miR146a to skin tissue and other organs

The in vivo stability and tissue distribution of LNPs are critical determinants of their therapeutic performance, particularly for sustained delivery applications. Accordingly, we first examined LNP persistence and tissue localization using a subcutaneous dorsal skin model in wild-type mice. Cy5.5-labeled LNPs (Cy5.5-LNPs) were injected into the dorsal skin and longitudinally imaged using an IVIS Spectrum system. Sustained epifluorescence was observed at the injection site from day 0 through day 7, with a marked reduction by day 14 (Figure 8A). Quantitative analysis revealed that total radiant efficiency decreased to approximately 10% of the initial signal by day 14 (Figure 8B). Ex vivo imaging of major organs (skin, dorsal muscle, liver, lungs, heart, kidneys, and spleen) harvested on day 14 demonstrated no detectable accumulation of Cy5.5-LNPs outside the injection site (Figure 8C). Quantification confirmed that fluorescence was localized predominantly within the dermal tissue, with negligible signal detected in other organs (Figure 8D).

**Figure 8.**
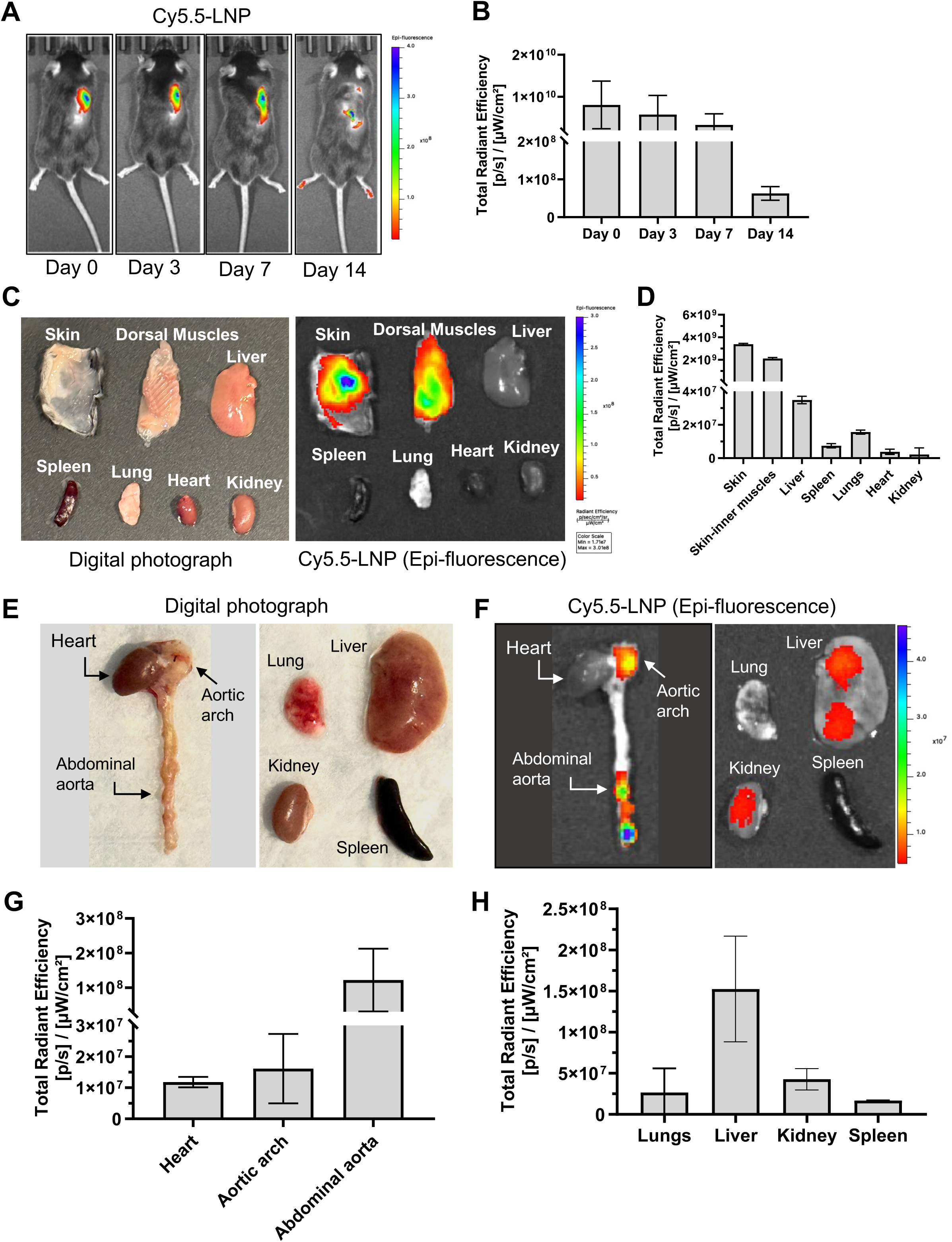
IVIS-based analysis of subcutaneous retention and systemic biodistribution of LNPs in vivo. In vivo uptake and biodistribution of Cy5.5-LNP in a murine subcutaneous skin model were evaluated using IVIS imaging from day 0 to day 14 **(A)**, with total radiant efficiency quantified **(B)**. Leaching and migration of Cy5.5-LNP to different organs – including skin, liver, dorsal muscle, spleen, lungs, heart, and kidneys - were checked **(C)**, and total radiant efficiency was quantified by IVIS analysis **(D)**. In addition, systemic migration and accumulation of Cy5.5-LNPs were assessed in hyperlipidemic ApoE⁻/⁻ mice. Representative digital images of major organs are shown **(E)**, and fluorescence intensity was analyzed by IVIS to determine total radiant efficiency **(F-H)**. All experiments were performed in triplicates, and data are presented as the mean ± standard error of the mean.

Systemic biodistribution of LNPs was next assessed following intravenous administration of Cy5.5-LNPs in hyperlipidemic ApoE⁻/⁻ mice. Representative digital images of excised major organs (heart, aorta, lung, liver, kidney, and spleen) are shown in Figure 8E. Epifluorescence imaging revealed preferential accumulation of Cy5.5-LNPs in the aortic arch and abdominal aorta, while minimal signal was detected in the heart (Figure 8F). Quantitative analysis of total radiant efficiency demonstrated significantly higher accumulation in the abdominal aorta compared to the aortic arch and heart (Figure 8G). Consistent with expected nanoparticle clearance pathways, the liver exhibited the highest overall fluorescence intensity among all organs examined (Figure 8F), which was confirmed by radiant efficiency quantification (Figure 8H).

Encouraged by above findings, we further evaluated the stability and tissue distribution of miR146a-loaded nanoparticles using LNP-Cy3-Scr-miR146a in the subcutaneous skin model. Following dorsal skin injection, LNP-Cy3-Scr-miR146a was imaged immediately and over time using IVIS. As shown in Figure 9A, robust epifluorescence was detected at 0 h, 3 h, and day 1 within a consistent spatial radius at the injection site. In contrast, mice injected with HEPES/Cy3-Scr-miR146a (naked miR) exhibited a rapid decline in fluorescence intensity beginning at 3 h, with progressive signal loss by day 7. Although epifluorescence from LNP-Cy3-Scr-miR146a also gradually decreased, it remained detectable through day 4 and was substantially higher than naked miR at corresponding time points. Quantitative analysis of total radiant efficiency confirmed significantly faster signal decay for HEPES/Cy3-Scr-miR146a compared with LNP-Cy3-Scr-miR146a (Figure 9B).

**Figure 9.**
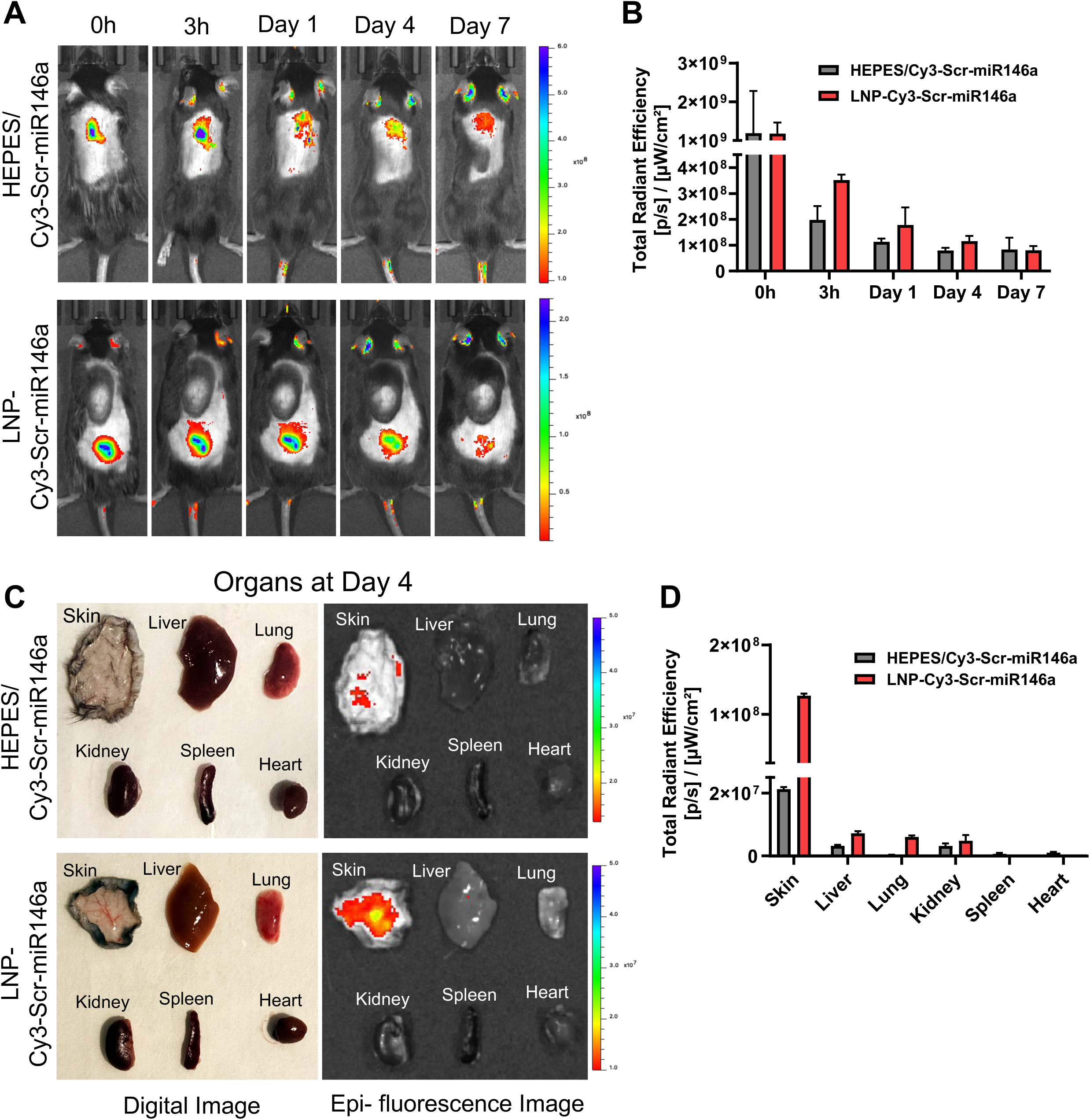
In vivo stability and tissue distribution of miR-146a-loaded LNPs. **(A)** In vivo skin tissue uptake and distribution of LNP-Cy3-Scr-miR146a and control HEPES/Cy3-Scr-miR146a were investigated by IVIS imaging at 0 h, 3 h, day 1, day 4, and day 7. **(B)** Quantification of total radiant efficiency for LNP-Cy3-Scr-miR146a and HEPES/Cy3-Scr-miR146a. **(C)** On day 4 leaching and migration of LNP-Cy3-Scr-miR146a and HEPES-Cy3-Scr-miR146a to different organs – including skin, liver, lungs, kidneys, spleen, and heart - were evaluated. **(D)** Quantification of total radiant efficiency in different organs as analyzed by IVIS imaging. Experiments were performed in triplicate, and data are presented as the mean ± standard error of the mean.

Tissue distribution of both formulations was further examined on day 4 post-injection by ex vivo imaging of skin and major organs, including liver, lung, kidney, spleen, and heart (Figure 9C). No evidence of systemic migration or off-target accumulation was observed for either formulation, with fluorescence signals confined predominantly to the injection site. Quantitative analysis of total radiant efficiency confirmed significantly higher signal intensity in the skin compared with other organs for both HEPES/Cy3-Scr-miR146a and LNP-Cy3-Scr-miR146a, with the latter exhibiting superior retention (Figure 9D).

Collectively, these results demonstrate that the engineered LNP-miR146a exhibit prolonged in vivo stability, minimal off-target distribution, and sustained retention at the site of administration, supporting their suitability as effective delivery vehicles for localized and targeted therapeutic applications.

### Delivery of miR-146a by LNPs prevents foreign body giant cell formation in vitro and attenuates inflammatory gene expression in vivo in an implantation model

After establishing the in vivo stability and tissue distribution of LNP-miR146a, its therapeutic efficacy was evaluated using both in vitro and in vivo models. First, miR-146a overexpression was assessed in BMDMs 24 h after treatment with LNP-miR-146a and LNP-Scr-miR146a. A robust ∼400-fold increase in miR-146a expression was observed compared with cells treated with LNP-Scr-miR146a (Figure 10A). To confirm the functional activity of the delivered miR-146a, expression of tumor necrosis factor receptor–associated factor 6 (TRAF6), a validated target of miR-146a, was examined in the same samples. A ∼30% reduction in TRAF6 gene expression was detected in LNP-miR146a-treated cells relative to the scrambled control (Figure 10B), confirming effective intracellular delivery and target engagement.

**Figure 10.**
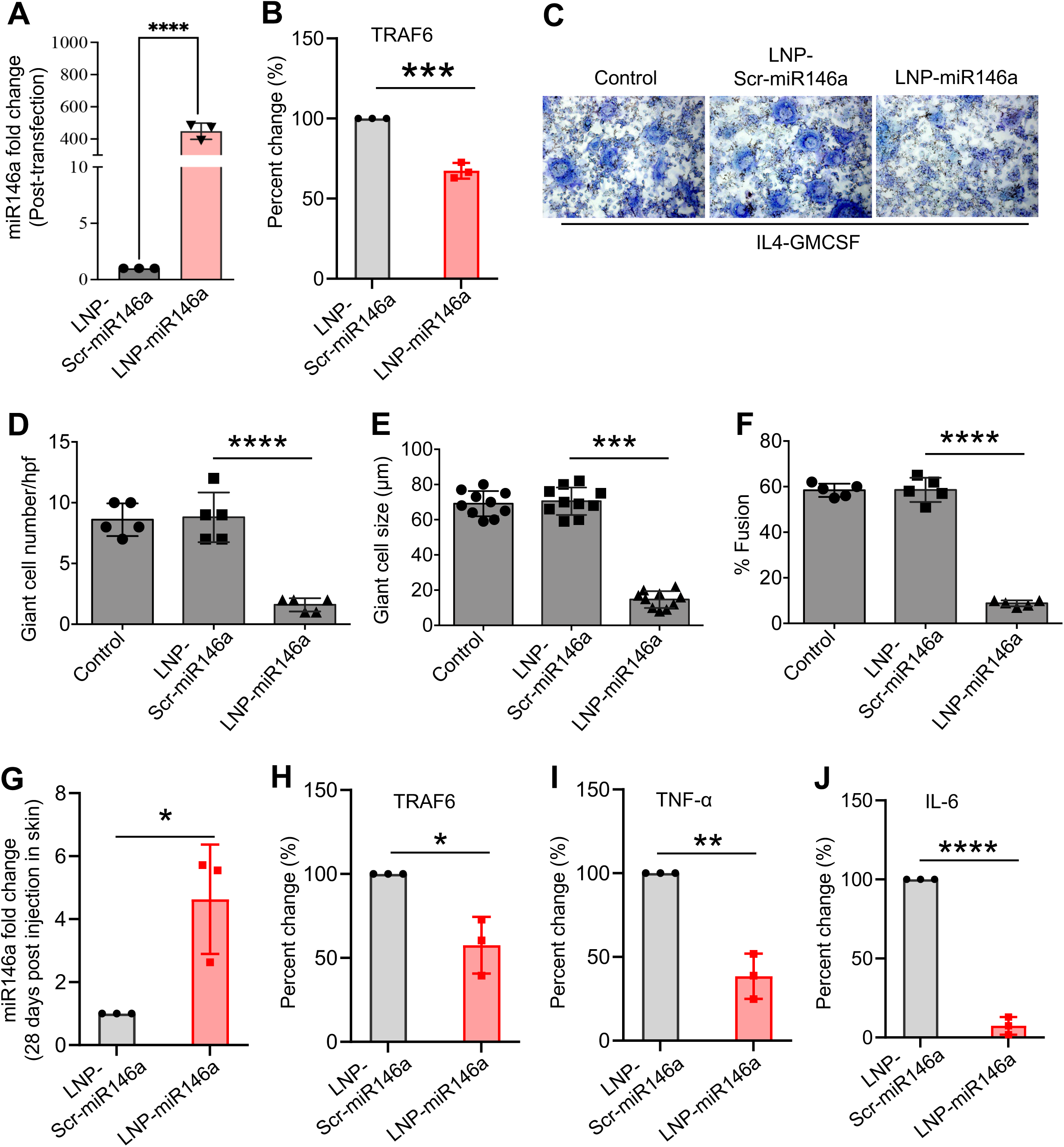
LNP-mediated miR-146a delivery attenuates foreign body giant cell formation and in vivo inflammation. The therapeutic potential of miR-146a-loaded LNPs, compared with the control LNP-Scr-miR146a, was investigated in BMDMs by assessing miR-146a overexpression **(A)**, attenuation of TRAF6 expression **(B)**, and suppression of foreign body giant cell formation as quantified by giant cells number, size, and fusion efficiency **(C-F)**. In vivo, LNP-miR146a-mediated overexpression of miR-146a and its functional activity in skin tissues were evaluated by measuring miR-146a and TRAF6 expression in a murine subcutaneous implantation model 28 days after implantation (**G, H**). Suppression of inflammation by LNP-miR146a administration was further quantified by assessing expression level of IL-6 and TNF-α in implanted skin tissues **(I, J)**. All experiment were performed in triplicate, and statistical significance was determined using ANOVA or unpaired t-test and is denoted as *p<0.05, **p<0.01, ***p<0.001, ****p<0.0001.

Based on these findings, we next investigated whether LNP-miR146a could inhibit FBGC formation, a hallmark of the foreign body response to biomaterial implantation (20,45,46). BMDMs pretreated with LNP-miR146a or LNP-Scr-miR146a were induced by IL-4 plus GMCSF to undergo fusion, and multinucleated giant cell formation was assessed by microscopy. Representative images demonstrated a marked reduction in FBGC formation in LNP-miR146a-treated cells compared with untreated controls and LNP-Scr-miR146a-treated cells (Figure 10C). Quantitative analysis revealed significant decreases in giant cell number, cell size, and fusion index in the LNP-miR146a group (Figure 10D-F). These data indicate that pretreatment with LNP-miR146a effectively suppresses macrophage fusion and FBGC formation, a critical process in FBR, a chronic inflammatory response to implantation.

To extend these findings in vivo, miR-146a expression was evaluated in murine skin tissue surrounding implanted cellulose ester membranes 28 days after subcutaneous (s.c) administration of LNP-miR146a or LNP-Scr-miR146a at days 0 and 14. A ∼4-fold increase in miR-146a expression was detected in LNP-miR146a-treated tissues compared with controls (Figure 10G). To assess downstream anti-inflammatory effects, the expression of TRAF6, interleukin-6 (IL-6), and tumor necrosis factor-α (TNF-α) was analyzed. All three inflammatory markers were significantly downregulated in LNP-miR146a-treated skin relative to LNP-Scr-miR146a (Figure 10H-J), demonstrating functional miR-146a delivery and sustained anti-inflammatory activity in a s.c implantation model.

### Discussions

The clinical translation of microRNA therapeutics has long been hindered by the inherent fragility of nucleic acids in the systemic circulation. Our study addresses these challenges by developing a robust, four-component LNP platform specifically optimized for the delivery of miR-146a, a critical "molecular brake" of the inflammatory response. The therapeutic efficacy of our platform stems from the strategic selection of lipid components. By utilizing a lipid injection-based formulation, we achieved a uniform size distribution and exceptional stability across a broad pH range (2.5-8). This stability is paramount for surviving the acidified environment of the endo/lysosomal pathway. The inclusion of DOTAP provided the necessary cationic charge to condense miR-146a and protect it from enzymatic degradation, while PEG served as a crucial stabilizer to prevent nanoparticle aggregation and manage surface moiety (11–13). A distinctive feature of our formulation is the synergistic use of DOPE and linoleic acid. DOPE’s fusogenic properties, combined with the membrane-fluidizing effects of unsaturated linoleic acid, likely facilitated the efficient endosomal escape observed in our results, ensuring that the miR-146a cargo reached the cytosol to exert its biological function (11–13).

miRs have emerged as potent endogenous "master switches" that regulate complex gene networks governing macrophage activation, inflammation, and fibrosis (14–38). Within this regulatory landscape, our recently published studies identify miR-146a as a central "molecular brake" essential for limiting the FBR to implanted biomaterials (20). We have demonstrated a definitive inverse relationship between miR-146a expression and immune severity: while elevated levels significantly suppress macrophage accumulation, FBGC formation, and collagen deposition, the genetic deletion of miR-146a in murine models exacerbates these inflammatory hallmarks. Consequently, miR-146a gene therapy represents a promising strategy to mitigate the FBR and improve device integration (20). In the present study, we show that the delivery of miR-146a via a specialized LNP platform effectively prevents FBGC formation *in vitro* and significantly attenuates inflammatory gene expression *in vivo* within a murine implantation model.

While our above-stated results are promising, several limitations must be acknowledged: i)while we observed robust tissue uptake and short-term efficacy, the long-term clearance rates and potential for off-target accumulation in other organs over extended periods remain to be fully elucidated; ii) our *in vivo* validation was conducted in a murine implantation model. Given the physiological differences in immune system complexity between mice and humans, further studies in large animal models are required to confirm the translatability of these findings; and iii) although the current LNP formulation showed high loading efficiency, we have not yet established the minimum effective dose required to maintain the "molecular brake" effect over the entire lifespan of a long-term implant, which may necessitate different release kinetics.

Collectively, our current results suggest this LNP-miR146a platform as a stable, efficient, and translatable approach for mitigating inflammation and addressing biomaterial implantation-associated inflammatory and FBR reactions. The observed potent anti-inflammatory activity in both primary cells and murine models, coupled with excellent biocompatibility, suggests that this LNP system overcomes the traditional barriers of toxicity and delivery inefficiency. The scalability of the lipid injection method further enhances the translatable potential of this platform for large-scale clinical applications.

## Acknowledgements

This work was supported by an NIH (R01EB024556) grant to Shaik O. Rahaman. We acknowledge the DLAR Imaging Core Facility, located at A.J. Clark Hall at the University of Maryland, College Park for providing imaging equipment assistance and consultation.

## Authors contributions

SOR and MIK conceived the study, designed the experiments, analyzed the data, and wrote the manuscript. MIK and KRS performed the experiments. SOR edited the manuscript. All authors reviewed and approved the final content of the manuscript.

## Data availability

All data generated and used during this study are included in this article.

## References

1. Ji, X., Meng, Y., Wang, Q., Tong, T., Liu, Z., Lin, J., Li, B., Wei, Y., You, X., Lei, Y. and Song, M. Cysteine-based redox-responsive nanoparticles for fibroblast-targeted drug delivery in the treatment of myocardial infarction. ACS nano. 2023;17(6):5421–5434.

2. de Castro Leao, M., Pohlmann, A.R., Alves, A.D.C.S., Farsky, S.H.P., Uchiyama, M.K., Araki, K., Sandri, S., Guterres, S.S. and Castro, I.A. Docosahexaenoic acid nanoencapsulated with anti-PECAM-1 as co-therapy for atherosclerosis regression. Eur. J. Pharm. Biopharm. 2021;159:99–107.

3. Khan, M.I., Paul, P., Behera, S.K., Jena, B., Tripathy, S.K., Lundborg, C.S. and Mishra, A. To decipher the antibacterial mechanism and promotion of wound healing activity by hydrogels embedded with biogenic Ag@ ZnO core-shell nanocomposites. Chem. Eng. J. 2021;417:128025.

4. Huang, Y., Guo, X., Wu, Y., Chen, X., Feng, L., Xie, N. and Shen, G. Nanotechnology’s frontier in combatting infectious and inflammatory diseases: prevention and treatment. Signal Transduct. Target. Ther. 2024;9(1):34.

5. Polack, F.P., Thomas, S.J., Kitchin, N., Absalon, J., Gurtman, A., Lockhart, S., Perez, J.L., Perez Marc, G., Moreira, E.D., Zerbini, C. and Bailey, R. Safety and efficacy of the BNT162b2 mRNA Covid-19 vaccine. N. Engl. J. Med. 2020;383(27):2603–2615.

6. Baden, L.R., El Sahly, H.M., Essink, B., Kotloff, K., Frey, S., Novak, R., Diemert, D., Spector, S.A., Rouphael, N., Creech, C.B. and McGettigan, J. Efficacy and safety of the mRNA-1273 SARS-CoV-2 vaccine. N. Engl. J. Med. 2021;384(5):403–416.

7. Xiong S, and Liu C. Breaking the PEG barrier to boost mRNA-LNP therapeutics. Nat. Rev. Mater. 2025;10(11):799–800.

8. Xu, S., Hu, Z., Song, F., Xu, Y. and Han, X. Lipid Nanoparticles: Composition, Formulation and Application. Mol. Ther. Methods Clin. Dev. 2025;33(2):101463.

9. Jung, H.N., Lee, S.Y., Lee, S., Youn, H. and Im, H.J. Lipid nanoparticles for delivery of RNA therapeutics: Current status and the role of in vivo imaging. Theranostics. 2022;12(17):7509–7531.

10. Abaza, T., Mohamed, E.E. and Zaky, M.Y. Lipid nanoparticles: a promising tool for nucleic acid delivery in cancer immunotherapy. Med. Oncol. 2025;42(9):409.

11. Miyasaki, K., Han, S., Carton, O., Kandell, R.M., Gunn, J. and Kwon, E.J. Formulation methods for peptide-modified lipid nanoparticles. J. Control Release. 2025;385:114030.

12. Albertsen, C.H., Kulkarni, J.A., Witzigmann, D., Lind, M., Petersson, K. and Simonsen, J.B. The role of lipid components in lipid nanoparticles for vaccines and gene therapy. Adv. Drug. Deliv. Rev. 2022;188:114416.

13. Wang, J., Chen, R., Xie, Y., Qin, X., Zhou, Y. and Xu, C. Endo/Lysosomal-Escapable Lipid Nanoparticle Platforms for Enhancing mRNA Delivery in Cancer Therapy. Pharmaceutics. 2025;17(7):803.

14. Bartel, D.P. MicroRNAs: target recognition and regulatory functions. Cell. 2009;136(2):215–33.

15. Lindsay, M.A. microRNAs and the immune response. Trends Immunol. 2008;29(7):343–51.

16. Sen, C.K. MicroRNAs as new maestro conducting the expanding symphony orchestra of regenerative and reparative medicine. Physiol. Genomics. 2011;43(10):517–20.

17. Marques-Rocha, J.L., Samblas. M., Milagro, F.I., Bressan, J., Martinez, J.A., Marti, A. Noncoding RNAs, cytokines, and inflammation-related diseases. FASEB J. 2015;29(9):3595–611.

18. Wu, X.Q., Dai, Y., Yang, Y., Huang, C., Meng, X.M., Wu, B.M., and Li, J. Emerging role of microRNAs in regulating macrophage activation and polarization in immune response and inflammation. Immunology. 2016;148(3):237–48.

19. Sevignani, C., Calin, G. A., Siracusa, L.D., and Croce, C.M. Mammalian microRNAs: a small world for fine-tuning gene expression. Mamm. Genome. 2006;17:189–202.

20. Mahanty, M., Dutta, B., Ou, W., Zhu, X., Bromberg, J.S., He, X. and Rahaman, S.O. Macrophage microRNA-146a is a central regulator of the foreign body response to biomaterial implants. Biomaterials. 2025;314:122855.

21. McKiernan, P.J. and Greene, C.M. MicroRNA dysregulation in cystic fibrosis. Mediators Inflamm. 2015;2015:529642.

22. Pandit, K.V. and Milosevic, J. MicroRNA regulatory networks in idiopathic pulmonary fibrosis. Biochem. Cell Biol. 2015;93:129–137.

23. Salamo, O., Mortaz, E. and Mirsaeidi, M. Noncoding RNAs: new players in pulmonary medicine and sarcoidosis. Am. J. Respir. Cell Mol. Biol. 2018;58(2):147–156.

24. Pfeffer, S.R., Yang, C.H. and Pfeffer, L.M. The role of miR-21 in cancer. Drug Dev. Res. 2015;76(6):270–277.

25. Tahamtan, A., Teymoori-Rad, M., Nakstad, B. and Salimi, V. Anti-inflammatory microRNAs and their potential for inflammatory diseases treatment. Front. Immunol. 2018;9:1377.

26. Ying, S.Y., Chang, D.C. and Lin, S.L. The microRNA (miRNA): overview of the RNA genes that modulate gene function. Mol. Biotechnol. 2008;38(3):257–268.

27. Barnett, R.E., Conklin, D.J., Ryan, L., Keskey, R.C., Ramjee, V., Sepulveda, E.A., Srivastava, S., Bhatnagar, A. and Cheadle, W.G. Anti-inflammatory effects of miR-21 in the macrophage response to peritonitis. J. Leucoc. Biol. 2016;99(2):361–371.

28. Jayawardena, T.M., Egemnazarov, B., Finch, E.A., Zhang, L., Payne, J.A., Pandya, K., Zhang, Z., Rosenberg, P., Mirotsou, M. and Dzau, V.J. MicroRNA-mediated in vitro and in vivo direct reprogramming of cardiac fibroblasts to cardiomyocytes. Circ. Res. 2012;110(11):1465–1473.

29. Zheng, C.Z., Shu, Y.B., Luo, Y.L. and Luo, J. The role of miR-146a in modulating TRAF6-induced inflammation during lupus nephritis. Eur. Rev. Med. Pharmacol. Sci. 2017;21(5):1041–1048.

30. Saba, R., Sorensen, D.L., and Booth, S.A. MicroRNA-146a: A Dominant, Negative Regulator of the Innate Immune Response. Front. Immunol. 2014;5:578.

31. Bhatt, K., Lanting, L.L., Jia, Y., Yadav, S., Reddy, M.A., Magilnick, N., Boldin, M., and Natarajan, R. Anti-inflammatory role of microRNA-146a in the pathogenesis of diabetic nephropathy. J. Am. Soc. Nephrol. 2016;27(8):2277–88.

32. Javidan, A., Jiang, W., Okuyama, M., Thiagarajan, D., Yang, L., Moorleghen, J.J., Muniappan, L., and Subramanian, V. miR-146a Deficiency Accelerates Hepatic Inflammation Without Influencing Diet-induced Obesity in Mice. Sci. Rep. 2019;9(1):12626.

33. Su, Y.L., Wang, X., Mann, M., Adamus, T.P., Wang, D., Moreira, D.F., Zhang, Z., Ouyang, C., He, X., Zhang, B., Swiderski, P.M., Forman, S.J., Baltimore, D., Li, L., Marcucci, G., Boldin, M.P., and Kortylewski, M. Myeloid cell-targeted miR-146a mimic inhibits NF-kappaB-driven inflammation and leukemia progression in vivo. Blood. 2020;135(3):167–180.

34. Runtsch, M.C., Nelson, M.C., Lee, S.H., Voth, W., Alexander, M., Hu, R., Wallace, J., Petersen, C., Panic, V., Villanueva, C.J., Evason, K.J., Bauer, K.M., Mosbruger, T., Boudina, S., Bronner, M., Round, J.L., Drummond, M.J., and O’Connell, R.M. Anti-inflammatory microRNA-146a protects mice from diet-induced metabolic disease. PLoS Genet. 2019;15(2):e1007970.

35. Zou, Y., Cai, Y., Lu, D., Zhou, Y., Yao, Q., and Zhang S. MicroRNA-146a-5p attenuates liver fibrosis by suppressing profibrogenic effects of TGFβ1 and lipopolysaccharide. Cell Signal. 2017;39:1–8.

36. Jang, S.Y., Park, S.J., Chae, M.K., Lee, J.H., Lee, E.J., and Yoon, J.S. Role of microRNA-146a in regulation of fibrosis in orbital fibroblasts from patients with Graves’ orbitopathy. Br. J. Ophthalmol. 2018;102(3):407–414.

37. Du, J., Niu, X., Wang, Y., Kong, L., Wang, R., Zhang, Y., Zhao, S., and Nan, Y. MiR-146a-5p suppresses activation and proliferation of hepatic stellate cells in nonalcoholic fibrosing steatohepatitis through directly targeting Wnt1 and Wnt5a. Sci. Rep. 2015;5:16163.

38. Bobba, C.M., Fei, Q., Shukla, V., Lee, H., Patel, P., Putman, R.K., Spitzer, C., Tsai, M., Wewers, M.D., Lee, R.J., Christman, J.W., Ballinger, M.N., Ghadiali, S.N., and Englert, J.A. Nanoparticle delivery of microRNA-146a regulates mechanotransduction in lung macrophages and mitigates injury during mechanical ventilation. Nat. Commun. 2021;12(1):289.

39. Yu, B., Hsu, S.H., Zhou, C., Wang, X., Terp, M.C., Wu, Y., Teng, L., Mao, Y., Wang, F., Xue, W. and Jacob, S.T., 2012. Lipid nanoparticles for hepatic delivery of small interfering RNA. Biomaterials. 2012;33(25):5924–5934.

40. Kang, E. and Kortylewski, M. Lipid nanoparticle-mediated delivery of miRNA mimics to myeloid cells. In Inflammation and cancer. Methods. Mol. Biol. 2023;2691:337–350. (New York, NY: Springer US).

41. Schubert, M.A. and Muller-Goymann, C.C. Solvent injection as a new approach for manufacturing lipid nanoparticles–evaluation of the method and process parameters. Eur. J. Pharm. Biopharm. 2003;55(1):125–131.

42. Suri, K., Hosur, V., Panchakshari, R. and Amiji, M.M. A Multimodal Therapeutic Strategy for Inflammatory Bowel Disease Using MicroRNA-146a Mimic Encapsulated in Lipid Nanoparticles. Mol. Pharm. 2025;22(6):3132–3141.

43. Sharma, S., Mahanty, M., Rahaman, S.G., Mukherjee, P., Dutta, B., Khan, M.I., Sankaran, K.R., He, X., Kesavalu, L., Li, W. and Rahaman, S.O. Avocado-derived extracellular vesicles loaded with ginkgetin and berberine prevent inflammation and macrophage foam cell formation. J. Cell. Mol. Med. 2024;28(7):e18177.

44. Sharma, S., Goswami, R., Merth, M., Cohen, J., Lei, K.Y., Zhang, D.X. and Rahaman, S.O. TRPV4 ion channel is a novel regulator of dermal myofibroblast differentiation. Am. J. Physiol. Cell Physiol. 2017;312(5):C562–C572.

45. Goswami, R., Arya, R.K., Sharma, S., Dutta, B., Stamov, D.R., Zhu, X., and Rahaman, S.O. Mechanosensing by TRPV4 mediates stiffness-induced foreign body response and giant cell formation. Sc. Signal. 2021;14(707):eabd4077.

46. Mahanty, M., Ou, W., Zhu, X., Bromberg, J.S., He, X., and Rahaman, S.O. A Novel Core-Shell Hydrogel 3D Model for Studying Macrophage Mechanosensing and Foreign Body Giant Cell Formation. Adv. Healthc. Mater. 2025:e01614.

47. Gerlier, D., and Thomasset, N. Use of MTT colorimetric assay to measure cell activation. J. Immunol. Methods. 1986;94(1-2):57–63.

48. Khan, M.I., Behera, S.K., Paul, P., Das, B., Suar, M., Jayabalan, R., Fawcett, D., Poinern, G.E.J., Tripathy, S.K. and Mishra, A. Biogenic Au@ ZnO core–shell nanocomposites kill *Staphylococcus aureus* without provoking nuclear damage and cytotoxicity in mouse fibroblasts cells under hyperglycemic condition with enhanced wound healing proficiency. Med. Microbiol. Immunol. 2019;208(5):609–629.

